# *Cis* regulation within a cluster of viral microRNAs

**DOI:** 10.1101/2020.11.19.389551

**Authors:** Monika Vilimova, Maud Contrant, Ramy Randrianjafy, Philippe Dumas, Endrit Elbasani, Päivi Ojala, Sébastien Pfeffer, Aurélie Fender

## Abstract

MicroRNAs (miRNAs) are small regulatory RNAs involved in virtually all biological processes. Although many of them are co-expressed from clusters, little is known regarding the impact of this organization on the regulation of their accumulation. In this study, we set to decipher a regulatory mechanism controlling the expression of the ten clustered pre-miRNAs from Kaposi’s sarcoma associated herpesvirus (KSHV). We measured *in vitro* the efficiency of cleavage of each individual pre-miRNA by the Microprocessor and found that pre-miR-K1 and -K3 were the most efficiently cleaved pre-miRNAs. A mutational analysis showed that, in addition to producing mature miRNAs, they are also important for the optimal expression of the whole set of miRNAs. We showed that this feature depends on the presence of a canonical pre-miRNA at this location since we could functionally replace pre-miR-K1 by a heterologous pre-miRNA. Further *in vitro* processing analysis suggests that the two stem-loops act in *cis* and that the cluster is cleaved in a sequential manner. Finally, we exploited this characteristic of the cluster to inhibit the expression of the whole set of miRNAs by targeting the pre-miR-K1 with LNA-based antisense oligonucleotides in cells either expressing a synthetic construct or latently infected with KSHV.

## INTRODUCTION

Kaposi’s sarcoma herpes virus (KSHV) or Human herpes virus 8 is a gammaherpesvirus associated with cancers such as Kaposi’s sarcoma, B-lymphomas or the proliferative disorder Castelman disease. Its genome is a ∼165 kb dsDNA molecule that encodes more than 90 open reading frames (ORFs) as well as 25 mature microRNAs (miRNAs) (1). KSHV establishes lifelong persistent infection with a restricted expression of viral genes. However, a small percentage (<3%) of cells support lytic replication and under certain conditions KSHV can reactivate from latency to lytic replication. A dynamic balance between the latent and lytic phases of KSHV replication is critical to establish a successful virus infection, maintain latency, and is involved in pathogenic effects such as tumorigenesis (reviewed in (2)).

Interestingly, all KSHV precursor (pre-)miRNAs are expressed on the same polycistronic transcript, which is associated with latency (3–5). Ten of them (pre-miR-K1 to -K9 and miR-K11) are clustered within an intron of ∼4 kb between ORF71 (v-FLIP) and the kaposin genes, and are expressed under the control of a latent promoter (6). Pre-miR-K10 and pre-miR-K12 localize within the ORF and the 3’UTR of the kaposin gene, respectively, and are controlled by both latent and lytic promoters (6, 7).

KSHV miRNAs are able to regulate the expression of both viral and cellular genes that are essential to virus infection and associated diseases. Abundantly expressed during the latent phase, they directly participate in its maintenance, for instance by repressing directly, or indirectly through targeting of NF-kB pathway, the replication and transcription activator (RTA), which is crucial for viral reactivation (8–10). They also promote tumorigenesis by modulating apoptosis, angiogenesis or cell cycle (e.g. (11–14)). Finally, KSHV miRNAs also enhance immune evasion and viral pathogenesis by regulating host immune responses (e.g. (15–18). See also (19) for a recent review on KSHV miRNA functions.

Although we now know of numerous functions of KSHV miRNAs due to active research in the field, we still do not have a precise understanding of the regulation of expression of these key viral factors. In animals, miRNA biogenesis is a multi-step process including two maturations by RNase III enzymes. MiRNA genes are generally transcribed by RNA polymerase II as a long primary transcript (pri-miRNA) of several kilobases that can contain one or several miRNA precursor hairpins (pre-miRNA). First, the pri-miRNA is processed in the nucleus by the Microprocessor, comprising the RNAse III type enzyme Drosha and its cofactor DGCR8. After export into the cytoplasm, the pre-miRNA is further processed by another RNase III enzyme, Dicer, associated with TRBP. The final result is a duplex of miRNAs (5p and 3p) from which one of the strands is preferentially incorporated into an Argonaute protein to form the RISC complex, which can then be directed toward target mRNAs (reviewed in (20)). Alternative pathways of miRNA biogenesis exist such as Drosha-independent processing of mirtrons or Dicer-independent Ago2-dependent miR-451 cleavage (21–24).

About 25 to 40% of human miRNAs are found in clusters (25, 26). There are usually two to three miRNAs in a cluster. However, a few larger clusters were also described such as the conserved mammalian pri-miR-17∼92 that contains 6 members, or the imprinted C19MC that contains 46 tandemly-repeated pre-miRNA genes (27–29). Co-expression may be essential as it was shown that many clustered miRNAs regulate common biological processes as is the case for KSHV miRNAs (e.g. (18)). Even though clustered miRNAs are co-transcribed, the resulting mature miRNAs are found at different levels in the cell (e.g. (30, 31). This suggests that complex regulation events occur downstream of the transcription. Two independent studies from Zeng and Orom laboratories revealed the key importance of maturation by the Microprocessor to explain the global level of cellular miRNAs (32, 33). Maturation of pri-miRNAs by the Microprocessor is controlled by sequence and structural features of miRNA hairpin, defining the basal level of pre-miRNAs excised. In addition, protein cofactors may interact with specific motifs or structure of the stem-loop and thus modulate the Microprocessor activity (see (34–36). Recently, several studies demonstrated interdependent processing in the context of bicistronic pri-miRNA where an optimal miRNA hairpin assists the processing of a neighboring suboptimal one (37–42). Such interdependency has not yet been documented in the case of larger miRNA clusters.

Previously, we demonstrated the importance of RNA secondary structure of the long primary transcript containing the ten intronic miRNAs from KSHV (pri-miR-K10/12) for the accumulation of mature miRNAs (31). Here, we show *cis* regulation within this large viral miRNA cluster. We observed that the ten miRNA hairpins from the KSHV intronic cluster are processed *in vitro* by the Microprocessor with different efficiencies. Intriguingly, high processing levels of miR-K1 and miR-K3 hairpins were not consistent with the low level of accumulation of their mature miRNAs in infected cells (31), suggesting that these miRNA hairpins could serve other purposes than solely producing mature miRNAs. Indeed, specific deletion of pre-miR-K1 or pre-miR-K3 within the cluster significantly reduce the expression of the remaining miRNAs in the cell. Moreover, only the pre-miRNA feature is sufficient to support such regulatory mechanism since replacement of pre-miR-K1 by the heterologous pre-Let-7a-1 restores expression of clustered miRNAs to the wt level. Further experiments of *in vitro* processing assays using pri-miRNA fragments (mimicking cleavage of pre-miR-K1 or pre-miR-K3) suggest that regulation may occur before pre-miR-K1 and -K3 are cut by the Microprocessor or that processing of the cluster is sequential. Finally, we developed an antisense strategy based on LNA molecules to post-transcriptionally downregulate the expression of the whole KSHV miRNA cluster. Using an LNA targeting either the 5p or 3p of the pre-miR-K1 sequence, we managed to significantly reduce the levels of clustered KSHV miRNAs in cells transfected with a synthetic construct. We also showed that in KSHV-infected cells, the levels of neosynthesized miRNAs derived from the cluster dropped significantly upon LNA targeting of pre-miR-K1, indicating that this could be a useful strategy to block the entire cluster in infected cells.

## MATERIALS AND METHODS

### Cells and media

HEK293FT-rKSHV cells were generated by infecting HEK293FT cells with concentrated rKSHV.219 virus (43) in the presence of 8 µg/mL polybrene (Sigma) and by spinoculation (800 g for 30 min at room temperature). Puromycin selection was applied to select for rKSHV.219 infected cells. The virus stock used to infect HEK293FT cells was generated by treating iSLK.219 cells (44) with 1 μg/mL doxycycline and 1.35 mM sodium butyrate and collecting virus particles 48 h post-reactivation by ultracentrifugation.

Adherent HEK293Grip and HEK293FT-rKSHV cell-lines were cultured in a humidified 5% CO_2_ atmosphere at 37°C in DMEM medium containing 10% fetal calf serum (FCS). In addition, HEK293FT-rKSHV were grown with 2,5 µg/mL puromycin (for viral genome maintenance).

### RNA preparation

Total RNA was extracted from cells using TRIzol reagent (Invitrogen, Thermo Fisher Scientific).

Pre-miRNAs, wt and mutants of pri-miR-K10/12, derived from BCBL-1 cell line, were transcribed from PCR-generated DNA templates carrying a T7 promoter (see Table S1 for primers). *In vitro* RNA synthesis was done by T7 RNA polymerase (Ambion). Pre-miRNAs were purified on denaturant polyacrylamide gel and long pri-miR-K10/12 derived transcripts (up to ∼3 kb) were salt purified using Monarch® PCR and DNA cleanup kit (New England BioLabs). After acidic phenol extraction and ethanol precipitation, the RNAs were pelleted and recovered in MilliQ water.

### Northern blot analysis

RNAs were resolved on a 8% urea-acrylamide gel, transferred on a nylon membrane (Amersham Hybond-NX, GE-Healthcare Life Sciences), crosslinked to the membrane by chemical treatment at 60°C using 1-ethyl-3-[3-dimethylaminopropyl]carbodiimide hydrochloride (EDC) (Sigma) for 1 h 30 min. MiRNAs and pre-miRNAs were detected with specific 5’-^32^P labeled oligonucleotides (Table S1). The signals were quantified using a Fuji Bioimager FLA5100. miR-16 was probed as a loading control.

### *In vitro* Drosha miRNA processing assays

Drosha and DGCR8 were overexpressed in 10-cm Petri Dish Hek293Grip cells using pCK-Drosha-Flag and pCK-Flag-DGCR8. After 48 h, cells were washed with ice-cold PBS, centrifuged and pellet was resuspended in 120 µL ice-cold lysis buffer (20 mM Tris-HCl pH 8.0, 100 mM KCl, 0.2 mM EDTA, 0.5 mM DTT, 5% glycerol and mini-complete EDTA-free protease inhibitor (Roche)). The cell suspension was sonicated during 5 min at high amplitude, 30 sec on and 30 sec off using Bioruptor™ UCD-200 (Diagenode), centrifuged for 10 min at 10 000 g, 4°C, and the supernatant was used for *in vitro* processing assays.

500 or 1000 fmol of *in vitro* transcribed wt or mutant pri-miR-K10/12 RNAs were denatured 3 min at 95°C, cooled on ice 3 min and folded in 1x structure buffer provided by Ambion during 30 min at 37°C. Processing assays were performed in 30 µL containing 15 µL of total protein extract (10 µg/µL) or 15 µL of lysis buffer, 6.4 mM MgCl_2_, 30 U Ribolock (Thermo Scientific™, Thermo Fisher Scientific). Just after addition of 170 µL elution buffer (2% SDS, 0.3 M sodium acetate), reaction was terminated by acidic phenol extraction followed by ethanol precipitation with 5 µg glycogen. After resuspension in formamide loading buffer, cleavage products were analyzed by northern blot. For quantification, *in vitro* transcribed and gel purified pre-miRNAs and synthetic miRNA oligonucleotides (IDT) were loaded at increasing concentration (from 1.5 to 25 fmol). A standard curve was generated by plotting the signal intensity against the amount of pre-miRNAs loaded and was used to calculate the absolute amount of pre-miRNAs produced by *in vitro* Drosha processing.

### Kinetic analysis

The experimental cleavage curves show different rates and different fractions of cleavage of each pre-miR. In addition, the experimental cleavage curves most often show a maximum followed by a slight decrease of the amount of pre-miR, which requires to introduce a secondary cleavage event of the newly formed pre-miR (either from a contaminant RNase, or from Dicer). The model in use is thus:

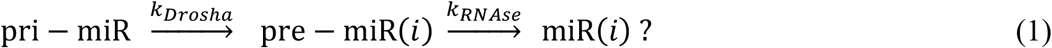

At first, it was attempted to model the rates of enzymatic cleavage (*k*_*Drosha*_ and *k*_*RNAse*_) according to Michaelis-Menten kinetics, but it turned out to be inefficient due to a very large correlation of the parameters *K*_*M*_ and *V*_*max*_. We finally used the following simplest possible mathematical model:

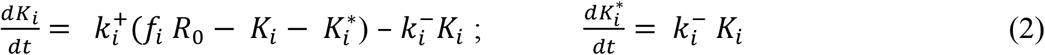

with *R*_0_ the total concentration of the pri-miRNA, *K*_*i*_ the time-dependent concentration of the *ith* pre-miR, *f*_*i*_ the fraction of *R*_0_ used by Drosha to produce *K*_*i*_, and 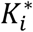 the time-dependent concentration of the secondary-cleavage product (miR(*i*) in equation (1)). Since Drosha act differently on the pri-miR to produce each pre-miR, it is necessary to replace *k*_*Drosha*_ with a particular cleavage rate 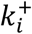 for each pre-miR and, similarly, it is necessary to replace *k*_*RNAse*_ with 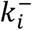 for each secondary-cleavage rate. This simple model, therefore, does not try in any way to differentiate situations wherein a particular pre-miR would be obtained either from the first cleavage of the full pri-miR, or from subsequent cleavage of a fragment of it. The previous linear differential equations (2) are readily integrated, which gives the cleaved fraction *Y*_*i*_ = *K*_*i*_ (*t*)/*R*_0_ necessary to fit the *i*^*th*^ experimental curve:

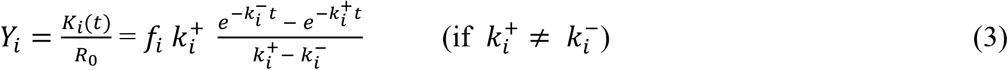

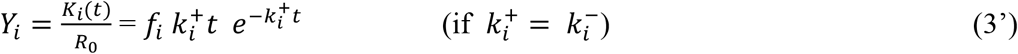

The results obtained with this model are in Figure 1 and Figure S2.

**Figure 1.**
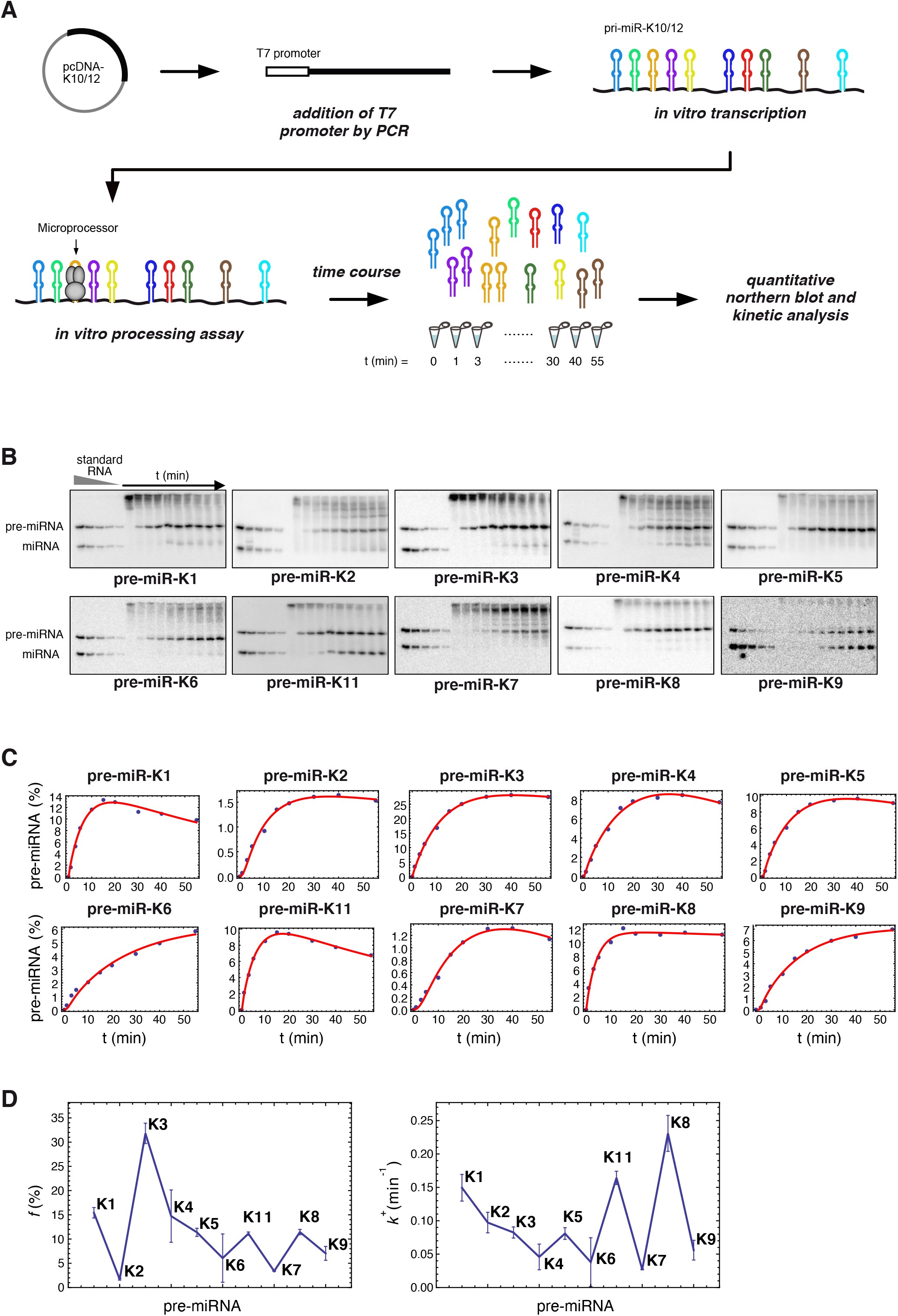
Kinetic analysis of KSHV clustered pre-miRNAs maturation *in vitro* by the Microprocessor. (A) Overview of *in vitro* processing assays starting with synthesis of DNA template containing a T7 promoter by PCR from the pcDNA-K10/12 plasmid and *in vitro* transcription to generate pri-miR-K10/12 containing the 10 KSHV clustered pre-miRNAs. *In vitro* processing assays was performed by incubating pri-miR-K10/12 with total protein extract of HEK293Grip cells overproducing Drosha and DGCR8. Pre-miRNAs production was monitored along the time and quantified by northern blot analysis. (B) Northern blot analysis of the time course of *in vitro* processing assays using 500 fmol of *in vitro* transcribed pri-miR-K10/12 and HEK293Grip cells total protein extract where Drosha and DGCR8 were overexpressed. *In vitro* transcribed pre-miRNAs and synthetic RNA oligonucleotides were loaded at increasing concentration as standards. (C) Cleavage curves were obtained after plotting pre-miRNA product, in percentage of initial pri-miR-K10/12 substrate, according to time. The fits were obtained with the model involving three free parameters per curve (compare with Figure S3 for the more stringent model with two free parameters per curve). (D) Processing efficiencies (left panel) and cleavage rate (right panel) were plotted in respect to miRNA hairpins showing variation among the clustered pre-miRNAs. The error bars come from standard procedures used to fit the experimental curves by minimizing the residuals between the experimental points and their theoretical estimates. Data are from Exp#1 (see Figure S2 for Exp#2).

### Mutagenesis and miRNA expression in human cells

Plasmid pcDNA-K10/12, derived from pcDNA5 (Invitrogen) and containing the wild type (wt) pri-miR-K10/12 (14) was mutated using the Phusion site-directed mutagenesis kit (Thermo Scientific) and transformation of *E. coli* DH5alpha strain. Positive clones were identified by sequencing (GATC Biotech, France). Four deletion mutants were designed: ΔK1, ΔK3, ΔK7 and ΔK9 where the respective pre-miRNA sequence, was deleted. ΔK1-Let7 corresponds to ΔK1 mutant where the pre-Let7a-1 was inserted in place of pre-miR-K1 (see Figure 2). Expression tests were conducted as follow: 2 µg of plasmids (wt or mutated) were used to transfect HEK293Grip cells in 6 well/plate. Total RNA was collected after 48h and miRNA expression was analyzed by northern blot, using 9 µg of total RNA and standard protocol. miR-16 was probed as a loading control and used for signal normalization.

**Figure 2.**
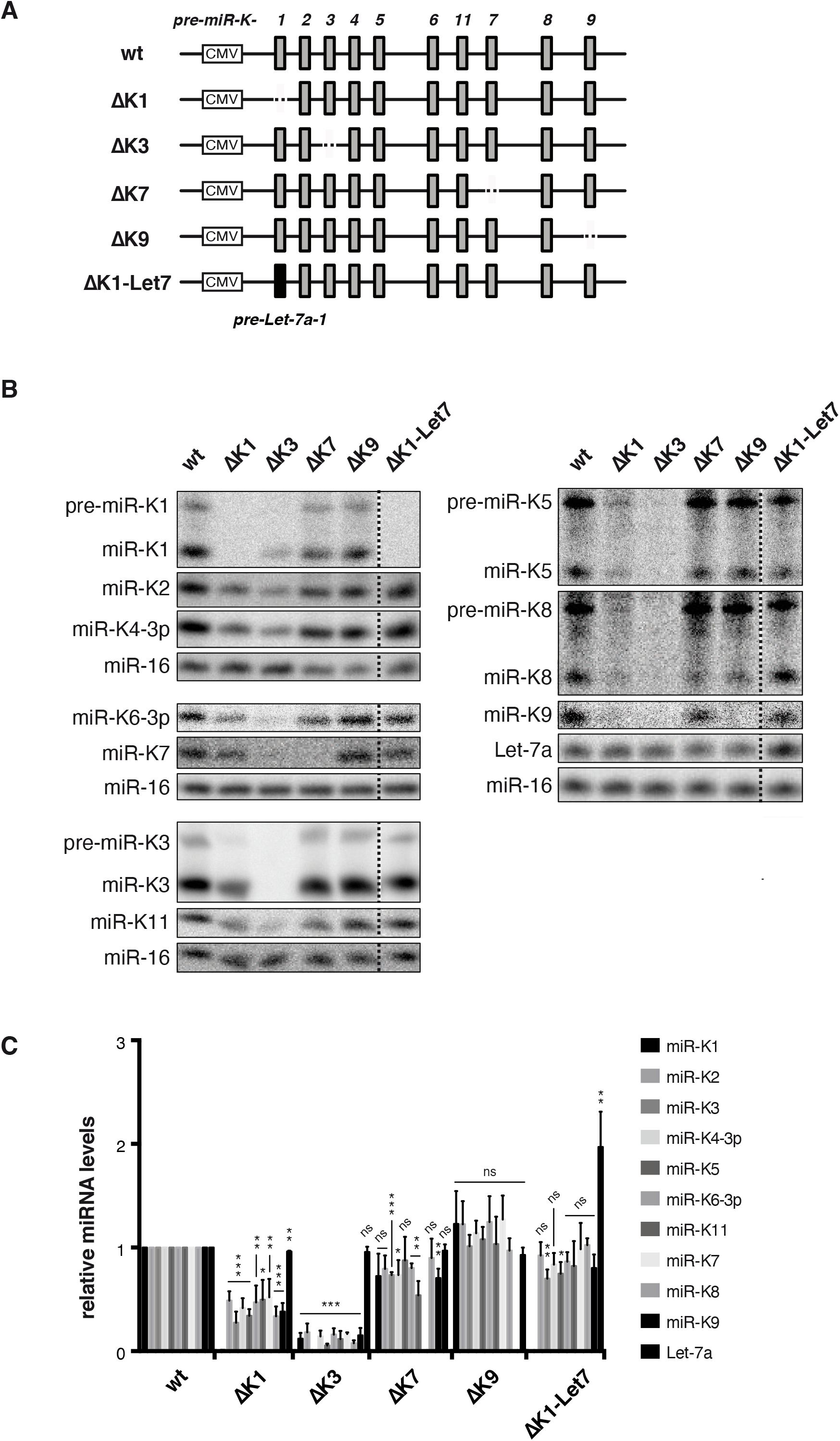
Mutational analysis reveals *cis* regulation within KSHV miRNA cluster. (A) Schematic view of pri-miR-K10/12 wt or mutant constructs used in the study. KSHV or hsa Let-7a-1 miRNA hairpins are represented by grey and black bars, respectively. Cytomegalovirus (CMV) promoter is shown. (B) Northern blot analysis of the accumulation of mature miRNAs, after overexpression of wt and mutant constructs in Hek293Grip cells (*n = 3*). MiR-16 was probed as a loading control. Dotted lines indicate where the blot was cut. (C) Histogram showing the relative expression of the clustered miRNAs from the mutated pri-miR-K10/12 constructs compared to the wt. Error bars were obtained from 3 independent experiments and p-values were obtained using unpaired t tests comparing wt versus mutant for each miRNA. ns: non-significant, *: p < 0.05, **: p <0.01, ***: p < 0.001.

### Antisense LNA treatments

HEK293Grip cells in 6 well/plate were transfected with 1 µg of wt pcDNA-K10/12 in combination with 20 nM LNA oligonucleotide (see Table S1 for sequence). Total RNA was collected after 48h and miRNA expression was analyzed by northern blot, using 10 µg of total RNA and standard protocol. miR-16 was probed as a loading control and used for signal normalization.

HEK293FT-rKSHV were seeded in 12 well/plates. When approximately 50-60% confluent, they were transfected with 20 nM LNA by using Lipofectamine 2000 reagent (Invitrogen). On the next day, they were detached by 50 µl of 0,05% Trypsin-EDTA (Gibco), resuspended in fresh medium and half of the cells were transferred into a new well and allowed to seed. On the next day they were transfected again with 20 nM LNA. They were collected by direct lysis in Trizol reagent (Ambion) one day after the second transfection.

### 4sU metabolic labeling, neosynthesized RNA pull-down and RT-qPCR analysis

The experimental procedure was adapted from the protocol developed by the Nicassio laboratory (45). One day prior to LNA transfection, 5 million HEK293FT-rKSHV cells were seeded in a 10 cm culture dish. 50nM LNA were transfected using Lipofectamine 2000 (Invitrogen) in total culture volume of 10 ml. After 24 h, culture medium was collected, filtered (45 µm) and 4sU was added to reach final concentration of 300 µM. The medium was then transferred back to the cells which were allowed to incorporate the 4sU during 3 h (incubation at 37°C, 5% CO_2_, in dark). After that, cells were detached in ice-cold PBS and collected by centrifugation. RNA was extracted by TRIzol reagent (Invitrogen) by using 3 ml of the reagent per dish.

70 µg of total RNA were biotinylated by incubation with 160 µl of EZ-Link HPDP-Biotin (Thermo Scientific, 1 mg/ml in DMF), and biotinylation buffer (final concentration 10 mM Tris pH 7.4, 1 mM EDTA) in total volume of 490 µl during 2 h at 25°C. Following PCI (Roth) extraction and isopropanol precipitation, the RNA was washed with EtOH 75% and dissolved in 80 µl of RNase-free water. Biotinylated RNA was then pulled-down on Dynabeads MyOne Streptavidin T1 (Invitrogen) by using 80 µl of beads per condition. The beads were first washed twice in buffer A (80 µl of 100 mM NaOH, 50 mM NaCl), then once in buffer B (100 mM NaCl) and finally resuspended in 160 µl of buffer C (2 M NaCl, 10 mM Tris pH 7.5, 1 mM EDTA, 0.1% Tween-20). The volume of RNA was increased to 160 µl and added to the beads. After 15 min of rotation on a wheel at room temperature, the beads with captured RNA were washed three times in 320 µl of buffer D (1 M NaCl, 5 mM Tris pH 7.5, 0.5 mM EDTA, 0.05% Tween-20). The RNA was then eluted with 160 µl of elution buffer (10 mM EDTA in 95% formamide) by heating at 65°C for 10 min. Trizol-LS (Invitrogen) and chloroform were used for eluted RNA extraction and after addition of 1.5 V of EtOH 100% to the aqueous phase, RNA was recovered on miRNeasy columns (Qiagen) in final volume of 30 µl.

3 µl of purified RNA were reverse-transcribed by using TaqMan™ MicroRNA Reverse Transcription Kit (Applied Biosystems) and a pool of eight specific stem-loop primers (miR-K1, miR-K3, miR-K4, miR-K11, miR-16, let-7a1, miR-92, U48, 0.5 µl each). RT reaction was then diluted twice and 1 µl used to perform qPCR in total volume of 10 µl, by using TaqMan™ Universal Master Mix II, no UNG (Applied Biosystems) and 0.5 µl of individual TaqMan miRNA assays (Applied Biosystem). qPCR was realized on CFX96 Touch Real-Time PCR Detection System (Biorad). Analysis of input RNA was performed in the same way on 100 ng of total RNA. In order to determine the amount of neosynthesized miRNAs relative to the input, we first calculated the enrichment of miRNA levels in the pull-down relative to the input after normalizing the data to Let-7. This ratio was then compared between the specific treatment (LNA @K1*) and the control treatment (LNA Ctrl), which was arbitrarily set to 1.

Primary transcript pri-miR-K10/12 was analyzed after prior treatment with DNase I (Invitrogen) or TURBO™ DNase (Invitrogen). 5 µl of purified or 1 µg of input RNA was treated and subsequently reverse-transcribed (using ¼ of reaction volume for non-reverse-transcribed control) with Superscript IV (ThermoFisher Scientific) according to the manufacturer protocol. cDNA was diluted twice before using 1 µl for quantitative PCR Maxima SYBR Green qPCR Master Mix (Thermo Scientific). The same approach than for mature miRNAs was used to determine the amount of neosynthesized pri-miRNAs except that the data were normalized to the CYC1 mRNA instead of Let-7.

RT-qPCR analysis of the primary transcript in HEK293Grip cells transfected with LNA was performed according to the same procedure, however by diluting the cDNA 10 times. miRNA expression in HEK293FT-rKSHV cells transfected with LNAs without metabolic labeling was measured similarly to the 4sU-samples, except for the reverse transcription step, which was performed individually for each RT stem-loop primer (0.5 µl) and without diluting the resulting cDNA. 100 ng of total RNA was used for each RT reaction. Primer sequences are indicated in the Table S1.

## RESULTS

### Clustered KSHV pre-miRNAs are processed *in vitro* by the Microprocessor with different efficiencies

In this work, we focused on the polycistronic feature of the KSHV intronic pri-miRNA containing ten miRNA hairpins (miR-K1 to miR-K9, and miR-K11), that we referred to as pri-miR-K10/12 (Figure S1). Our previous results showed that pri-miR-K10/12 adopts a well-organized 2D structure, composed of multiple hairpins with all miRNA sequences found in stem-loops. Interestingly, the secondary structural features of miRNA stem-loops correlate to some extent with the cellular abundance of mature miRNAs. Indeed, optimally folded stem-loops tend to lead to more abundant miRNAs. Moreover, we demonstrated that the structural context of miRNA hairpins within the primary transcript is important since swapping miRNA stem-loops or expressing miRNAs individually results in differential miRNA accumulation in cells (31).

To further understand the mechanism behind the regulation of expression of polycistronic KSHV miRNAs, we assessed the *in vitro* processing efficiency of the different pre-miRNAs within the cluster. To do so, we took advantage of *in vitro* processing assays using total extracts obtained from cells over-expressing Drosha and DGCR8 and *in vitro* transcribed pri-miR-K10/12 (∼3.2 kb) (Figure 1A). Accumulation levels of all pre-miRNAs from the cluster were analyzed at different time points using quantitative northern blot analysis (Figure 1B). Figure 1 and Figure S2 show the results obtained from two independent experiments (Exp#1 and Exp#2).

In the conditions used, all pre-miRNAs were produced from the unique pri-miR-K10/12 substrate. Pre-miRNAs were the major products, and, in some cases, we also observed accumulation of mature miRNAs due to residual Dicer activity in the total protein extract. Experimental cleavage curves were fitted using a simple kinetic model with three free parameters per curve (Figure 1C). This led to excellent agreement compared to a more stringent model with only two free parameters per curve (see Material and Methods and Supplemental Method). The numerical results are shown in Table S2. Interestingly, we noticed that the sum of all pre-miRNAs is (114 ± 8) % and (110 ± 17) % for Exp#1 and Exp#2, respectively, which is 10-fold lower that the maximum possible value of 1000 % if pri-miRNA gave rise to all ten possible pre-miRNAs. Since these two sums are not significantly different from 100 %, this suggests that each pri-miR-K10/12 would be cleaved only once and would produce only one particular pre-miRNA. However, we set our experimental conditions such that the initial concentration of substrate will be high enough to ensure that we measured the maximum rate of cleavage for each miRNA stem-loop. In that respect, the Microprocessor was probably saturated by the full-length pri-miR-K10/12 at the disadvantage of cleaved RNA fragments. As a consequence, we overlook the possibility to observe several rounds of cleavage.

Comparison of processing efficiencies *f*_*i*_ and of the cleavage rate constant 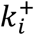 for each KSHV pre-miRNA showed important differences between pre-miRNAs as illustrated in Figure 1C and Figure S2C. Indeed, average accumulation levels vary from 1.4 % for pre-miR-K2 up to 36 % for pre-miR-K3 products (Table 1); cleavage rate constants ranged from 0.027 min^-1^ for pre-miR-K7 to 0.23 min^-1^ for pre-miR-K8 in Exp#1 (Figure 1 and Table S2). Kinetic parameters were not averaged due to a higher activity of the Microprocessor in Exp#1 *vs*. Exp#2. Drosha/DGCR8 concentration was estimated to be about three-fold greater in Exp#2 based on the relative values of cleavage rates in the two experiments. The agreement between the two experiments is rather good for the processing efficiencies *f*_*i*_ (correlation coefficient = 0.95) and lower for the kinetic constants of cleavage 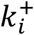 (correlation coefficient = 0.52) (figure S4). We can nevertheless conclude that there are significant differences of processing efficiencies between the different KSHV pre-miRNAs. To exclude that the observed differences might have arisen from various stability of the different pre-miRNAs and not their processing efficiencies, we assessed the stability of two pre-miRNAs that appeared well-processed in our experiment (pre-miR-K1 and -K8) and two pre-miRNAs that are less well-processed (pre-miR-K11 and - K7) (Figure S5). Briefly, *in vitro* transcribed pre-miRNAs were incubated in total cellular extract from HEK293Grip cells and pre-miRNAs stability was measured at various time points by northern blot analysis. Overall, we did not observe any striking difference in the decay rate between them, which could account for the variation measured in our *in vitro* cleavage assays. We previously determined hairpin optimality features for each KSHV pre-miRNA (31), which we decided to update in order to take into account novel features that have been published since (46, 47). In particular, we calculated the Shannon entropy for each miRNA hairpin as highlighted in Figure S6. The data obtained indicated that overall a lower Shannon entropy could be observed along the stem in well-processed miRNA hairpins, with the exception of stem-loop miR-K11 and miR-K9, as described previously (47). Table S3 summarizes the updated optimality features observed for KSHV pre-miRNAs. This allowed us to compare processing efficiencies *f*_*i*_ of miRNA hairpins with their hairpin optimality feature and their corresponding miRNA abundance in infected BCBL-1 cells (as previously determined (31)) (Table 1). MiRNA hairpins were ranked from the best to the worst substrates, which allowed us to define two groups, *i*.*e*. the well or moderately processed (*f*_*i*_ ≥ 10 %, miR-K1, -K3, -K4, -K8 and -K11) and the less efficiently processed (miR-K2, -K6, -K7 and -K9). The miR-K5 hairpin was not ranked since the results were too discordant between the two experiments. Overall, well processed miRNA hairpins corresponded to optimally folded stem-loops. However, with the exception of miR-K4, this did not correlate with the level of accumulation of mature miRNAs in infected cells (31). Thus, miR-K1 and -K3 are embedded within hairpins that are optimal and the best processed by the Microprocessor *in vitro* (16 and 36% respectively), whereas the level of mature miRNAs is quite low (∼3 and ∼8 %, respectively). Accordingly, they were defined as over-processed. On the opposite, miR-K11 hairpin is only processed at ∼10% although it is the most abundant miRNA in cells, representing ∼23 % of viral clustered miRNAs. This hairpin was thus defined as being sub-processed. The same held true for miR-K6 and -K7 hairpins. Of the remaining hairpins, only miR-K2, -K4 and -K9 showed a good correlation between their optimality feature, processing efficiencies and cellular abundance.

**Table 1.**
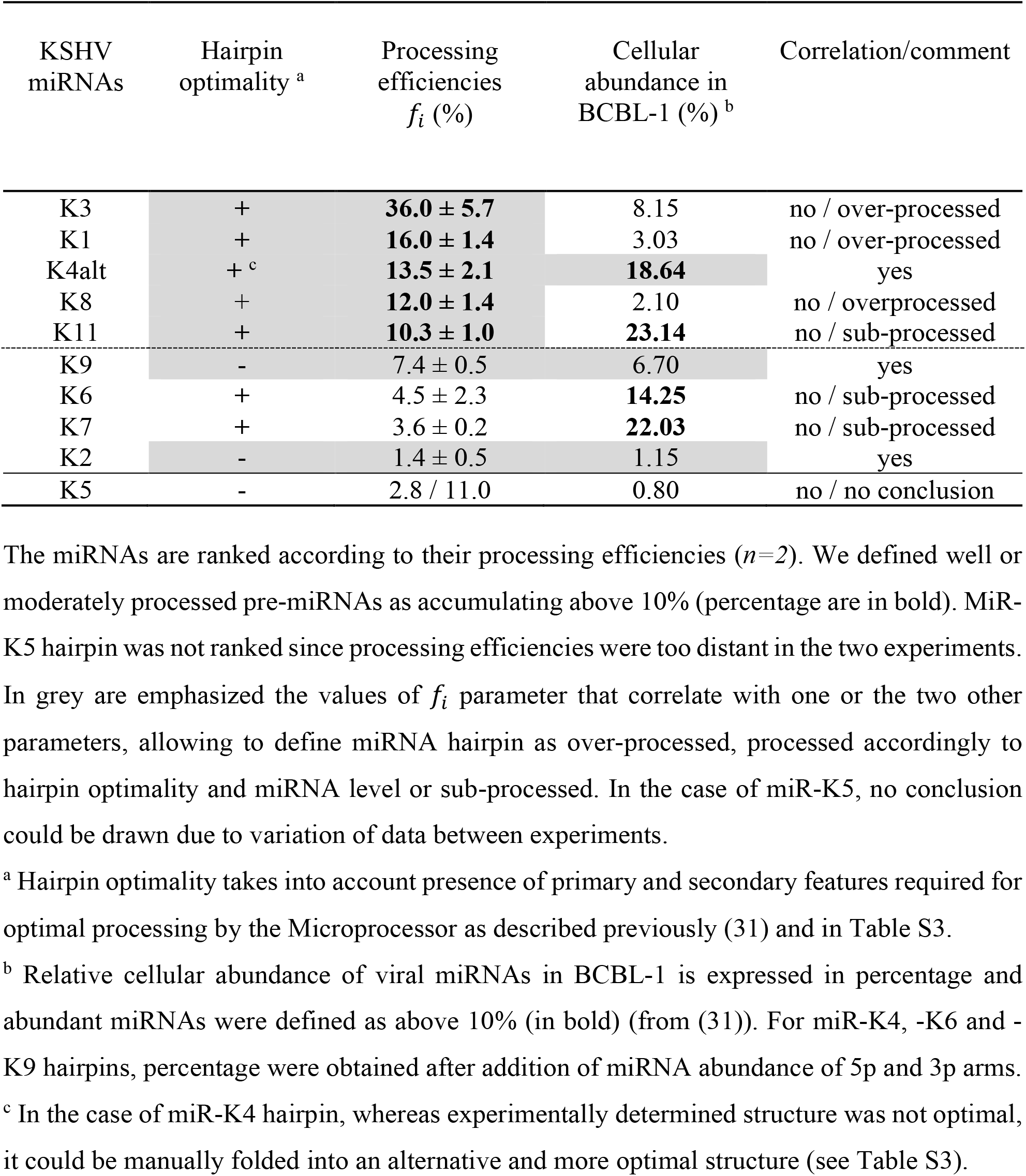
Correlation between KSHV miRNA hairpin processing efficiency with their hairpin optimality and cellular abundance of their corresponding mature miRNAs.

Altogether, our kinetic analysis therefore shows different processing efficiencies of KSHV miRNA hairpins within the polycistronic pri-miR-K10/12, which did not fully correlate with the fact that the miRNA hairpin was optimal or not or with the accumulation level of the respective mature miRNAs in infected cells. These discrepancies may reflect complex regulation of their biogenesis. Indeed, sub-processing of miRNA hairpins may be explained by the requirement of additional elements such as protein cofactors that may be absent or present at low levels in our *in vitro* assay. In addition, we also observed cases of over-processed miRNAs, such as miR-K1 and miR-K3, which suggests that processing by the Microprocessor may serve here another purpose than solely producing mature miRNAs.

### Deletion of pre-miR-K1 or pre-miR-K3 globally impairs the expression of the remaining miRNAs from the cluster

To further study the processing of the KSHV miRNA cluster, we used a plasmid allowing expression of the pri-miR-K10/12 sequence driven by a CMV promoter (14). Although it seems that there is a better accumulation of miRNAs at the 5’ extremity of the cluster, the wild type construct gives rise to all ten miRNAs and their relative expression level is close to what can be measured in latently infected BCBL1 cells (Figure S7).

To investigate the potential other role of miR-K1 and -K3 processing we generated mutant constructs in which we deleted individually pre-miR-K1 or pre-miR-K3 sequences within the polycistronic pri-miR-K10/12. Other miRNA sequences from the cluster were unchanged. We then assessed the impact of these deletions on the expression of the remaining clustered miRNAs in the cell. As negative controls, we deleted pre-miR-K7 and pre-miR-K9, that are located in the middle and at the 3’ end of the cluster, respectively, and are not well processed *in vitro* by the Microprocessor. The resulting mutants were named ΔK1, ΔK3, ΔK7 and ΔK9 (Figure 2A). These were expressed in HEK293Grip cells and the accumulation levels of all mature miRNAs from the cluster were assessed by northern blot analysis (Figure 2B and C). Interestingly, the expression of all miRNAs in the cluster was globally and drastically decreased compared to the wt construct in the ΔK1 and ΔK3 mutants, whereas it was only moderately or mostly unaffected in the ΔK7 and ΔK9 mutants. ΔK1 construct led to miRNA levels significantly reduced down to ∼28 % (3.6-fold) and ∼52 % (1.9-fold) for miR-K3 and miR-K7, respectively, when compared to the wt plasmid. All the miRNAs from the cluster were negatively affected, whatever the distance between pre-miR-K1 and the impacted miRNAs. For example, the farthest miR-K9 was even more impacted than the closest miR-K2 (∼38 % (2.6-fold) *vs* ∼49 % (2-fold), respectively) (Figure 2C). Deletion of pre-miR-K3 within the cluster was even more deleterious for the expression of the rest of the clustered miRNAs. Indeed, miRNA levels were decreased down to ∼6 % (16.7-fold) (miR-K5) or ∼18 % (5.6-fold) (miR-K2). In the case of ΔK7 mutant, a moderately negative impact was observed for some miRNAs with the most affected being miR-K11, which level was reduced down to ∼54 % (1.9-fold). In the case of miR-K11, this may be due to a local effect of pre-miR-K7 deletion on the folding and/or processing of the adjacent miR-K11 hairpin.

Altogether, our results suggest that deletion of pre-miR-K1 or -K3 within pri-miR-K10/12 globally and drastically impacts the expression of the remaining miRNAs from the cluster in cells. This global and severe impact is specific to pre-miR-K1 and pre-miR-K3 since it was not observed for two other pre-miRNAs in the cluster.

### Expression of clustered miRNAs can be rescued by replacing pre-miR-K1 by a heterologous pre-miRNA

The analysis of deletion mutants indicates that pre-miR-K1 and -K3 appear to be required for the optimal expression of KSHV clustered miRNAs. This may be due either to *(i)* their ability to recruit the Microprocessor, as stem-loop structures, or *(ii)* specific sequences that establish tertiary contacts within the cluster or that are recognized by protein cofactors.

To investigate which hypothesis should be favored, we inserted a heterologous pre-miRNA sequence in lieu of pre-miR-K1 into the ΔK1 mutant and measured the expression of the rest of the clustered miRNAs. We chose the pre-Let-7a-1 sequence since both its primary sequence and its secondary structure, especially in its apical loop, is very different from those of pre-miR-K1, therefore lowering the possibility to form the same tertiary contacts or to recruit the same cofactors. The resulting mutant ΔK1-Let7 construct was expressed in HEK293Grip cells and the expression of miRNAs was assessed by northern blot analysis (Figure 2B, C). Whereas pre-miR-K1 was as expected not produced from the mutant construct, Let-7a expression was increased ∼2-fold when compared to the endogenous expression, showing that Let-7a within the context of the cluster was fully recognized and processed by the Microprocessor. The level of accumulation of all the KSHV miRNAs was measured and compared to the wt construct. Interestingly, expression of all of them was restored almost to the wt level, showing that replacing pre-miR-K1 with a heterologous pre-miRNA is sufficient for optimal production of the other miRNAs in the cluster. In conclusion, our results suggest that a pre-miRNA structure at this position within the cluster is sufficient to optimize the expression of KSHV miRNAs from this construct.

### *In vitro*, mimicking initial cleavage of miR-K1 or miR-K3 hairpins impacts the processing of only few miRNA hairpins from the cluster

Pre-miR-K1 or pre-miR-K3 are necessary for the optimal expression of the other miRNAs from the cluster, and at least for pre-miR-K1, this is independent of primary sequence but rather relies on the presence of a miRNA stem-loop structure. According to that observation, we hypothesized that cleavage of pre-miR-K1 or pre-miR-K3 by the Microprocessor may help to favor the processing of the other miRNAs from the cluster. If true, an RNA mimicking the initial cut of one or the other of these two pre-miRNAs would lead to a better processing of the remaining pre-miRNAs.

To test this, we performed *in vitro* processing of two different *in vitro* transcribed RNA mimics, namely cut-K1 and cut-K3. Figure 3A gives a schematic view of the RNA molecules. Briefly, cut-K1 is composed of an RNA fragment containing pre-miR-K2 to pre-miR-K9 and including pre-miR-K11. Its 5’ end starts just downstream of the 3’ end of pre-miR-K1 3p arm, as it would be after Microprocessor cleavage. Cut-K3 RNA is the combination of two independent RNA fragments. One comprises pre-miR-K1 and pre-miR-K2 and its 3’ end finishes just upstream of the 5’ end of pre-miR-K3 5p arm. The second embeds pre-miR-K4 to pre-miR-K9, including pre-miR-K11, and its 5’ end starts just downstream of the 3’ end of pre-miR-K3 3p arm. These constructs were incubated with total protein extracts from cells over-expressing Drosha and DGCR8 and pre-miRNA products were analyzed by northern blot analysis (Figure 3B). We analyzed pre-miRNAs based on their proximity (pre-miR-K1, -K3 and -K4) or not (pre-miR-K6, -K7, -K11 and -K8) to the deleted pre-miRNAs and the fact that their levels obtained in our kinetic analysis were low compared to the corresponding mature miRNA levels measured in infected cells (miR-K6, -K7 and -K11). Pre-miR-K4 and -K8 were chosen as controls since they were both efficiently processed.

**Figure 3.**
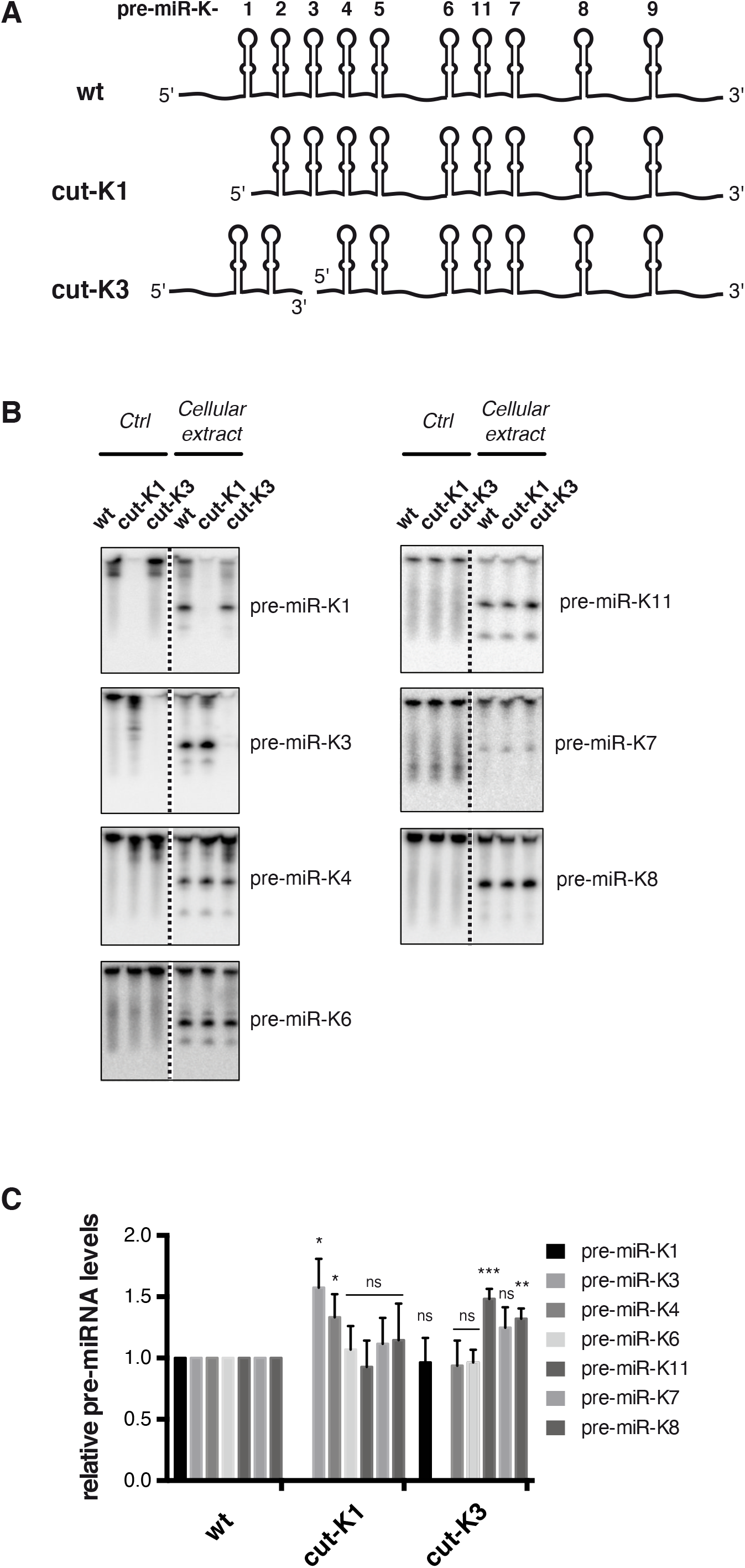
*In vitro* maturation assays using RNA mimicking miR-K1 and miR-K3 hairpins cleavage. (A) Schematic view of *in vitro* transcribed RNAs used in the study. Cut-K1 and cut-K3 RNAs mimic cleavage products by Drosha/DGCR8 of pre-miR-K1 and pre-miR-K3, respectively. As a result, cut-K3 is composed of 2 RNA fragments. (B) Northern blot analysis of pre-miRNAs produced after 45 min incubation of 1000 fmol of *in vitro* transcribed RNAs with HEK293Grip cells total protein extract where Drosha and DGCR8 were overexpressed (right part of blot) or lysis buffer (left part of blot). Dotted lines indicate where the blot was cut. (C) Histogram showing the relative level of pre-miRNAs compared to the wt. Error bars were obtained from 3 independent experiments and p-values were obtained using unpaired t tests comparing wt versus mutant for each pre-miRNA. ns: non-significant, *: p < 0.05, **: p <0.01, ***: p < 0.001.

Cut-K1 RNA gave rise to all the tested pre-miRNAs from the cluster, with the exception of pre-miR-K1 as expected. Whereas most of them are produced to similar levels as from the wt transcript, pre-miR-K3 and to a minor extent pre-miR-K4 accumulates significantly ∼1.6- and ∼1.3-fold more respectively when compared to wt condition.

RNA mimicking the cleavage of pre-miR-K3 led to significantly more pre-miR-K11 (∼1.5-fold) and to a milder extent more pre-miR-K8 (∼1.3-fold) whereas pre-miR-K1, -K4 and -K6 levels were unchanged. Pre-miR-K7 showed a small increase (∼1.25-fold) but this was not statistically significant. Whereas cut-K1 showed rather local effect, cut-K3 increased the processing levels of pre-miRNAs at long distances.

Altogether, our results show that initial processing of pre-miR-K1 or pre-miR-K3 does not dramatically improve *in vitro* the overall maturation by the Microprocessor of the other miRNAs within the cluster. On the contrary, it affects the processing of only few pre-miRNAs. These results may emphasize the necessity of miR-K1 or miR-K3 hairpins to be an integral part of the cluster to exert a *cis*-regulatory function and/or a sequential processing of the different pre-miRNAs.

### Blocking pre-miR-K1 cleavage by an antisense LNA oligonucleotide phenocopies its deletion

The use of antisense oligonucleotides has been described as an efficient approach to suppress miRNA function by sponging the mature miRNA (48). Interestingly, Hall et al. described that antisense LNA can also inhibit miRNA maturation steps (49). Indeed, an LNA oligonucleotide targeting the liver specific miR-122 also binds to pri-miR-122 and pre-miR-122, invading the stem-loop structure and hindering recognition by the Microprocessor and Dicer. This may account for ∼30% of the total inhibition of miR-122 activity. We therefore decided to use a similar strategy to block the processing of miR-K1 hairpin in order to downregulate the whole cluster. The LNA oligonucleotide that we used consists of 20 nt fully complementary to mature miR-K1-5p arm and contains 8 LNA residues in the middle part (from nt 8 to nt 15) (Table S1). Using a similar oligonucleotide, Gao and colleagues managed to efficiently suppress miR-K1 activity (9). In their study, they did not assess whether this was solely due to the sponging effect of mature miRNA, or whether this also decreased miRNA biogenesis.

The previously described construct containing the KSHV miRNAs cluster was transfected in HEK293Grip cells together with an LNA targeting miR-K1, namely LNA@K1, or a control LNA (Table S1). We then measured the levels of mature miRNAs from the entire cluster by northern blot analysis (Figure 4A, B). As expected, miR-K1 accumulation was strongly decreased (∼3.7-fold) upon treatment with LNA@K1 compared to treatment with the control LNA. Interestingly, the levels of all the other miRNAs within the cluster were also negatively affected (∼2.2-to almost 6-fold decrease) (Figure 4B). As a control, endogenous Let-7a was not affected, since its level was unchanged whatever the LNA treatment. Thus, inhibiting processing by antisense LNA targeting miR-K1 phenocopies the impact of ΔK1 mutant on the expression of the clustered miRNAs.

**Figure 4.**
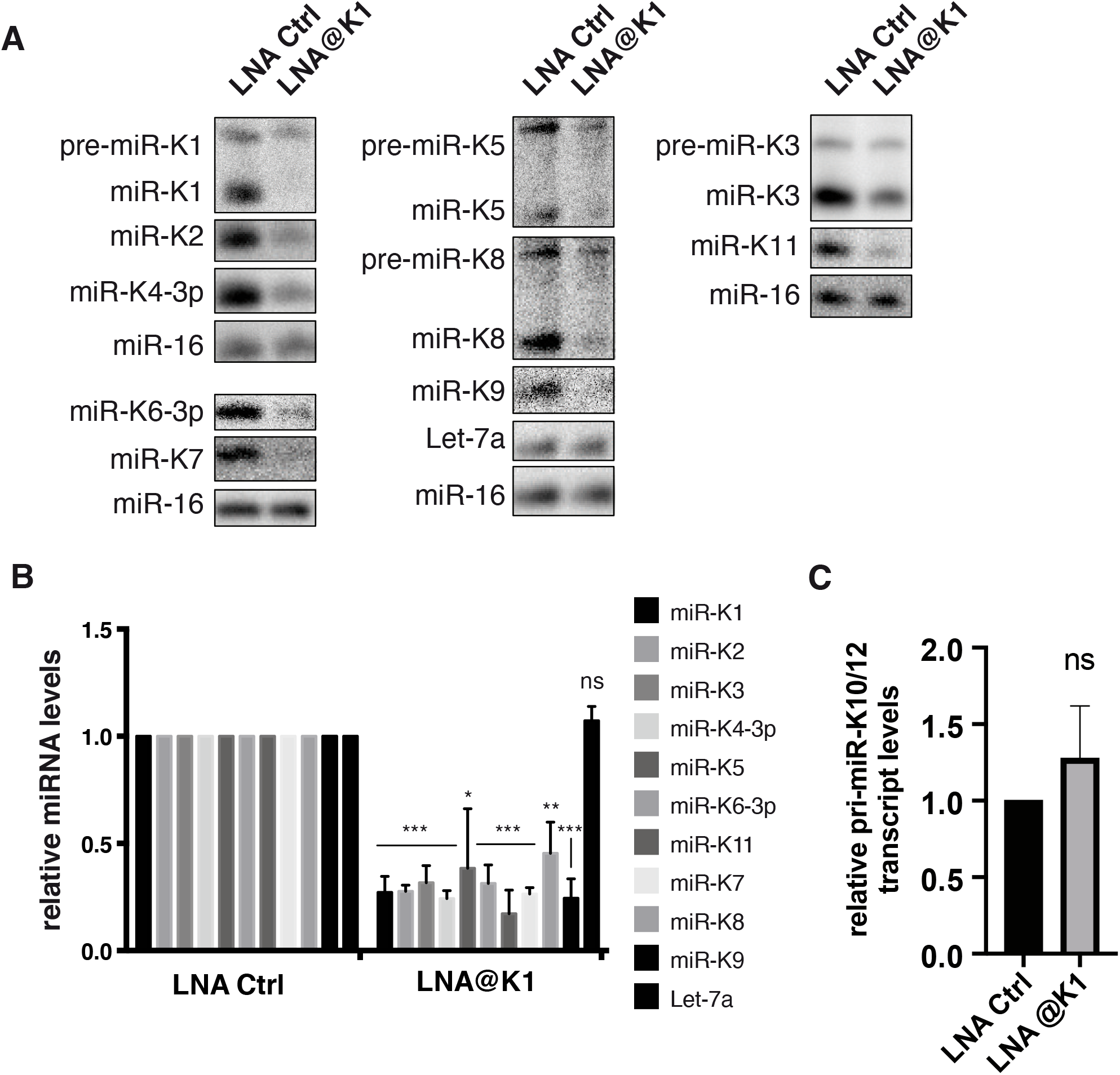
Antisense LNA targeting miR-K1 inhibits expression of other miRNAs from the cluster. (A) Northern blot analysis of the accumulation of mature miRNAs, after overexpression of wt plasmid in HEK293Grip cells during 48h (*n = 3*), with 20 nM control LNA (Ctrl LNA) or antisense LNA to miR-K1 (LNA@K1) treatment. Let-7a and miR-16 were probed as a control of miRNA expression and as a loading control, respectively. (B) Histogram showing the relative expression of the different miRNAs upon treatment with LNA@K1 compared to control LNA. Error bars were obtained from 3 independent experiments and p-values were obtained using unpaired t test with ns: non-significant, *: p < 0.05, **: p <0.01, ***: p < 0.001. (C) The expression of the KSHV miRNA primary transcript pri-miR-K10/12 was measured by RT-qPCR in total RNA samples from (B) and normalized to GAPDH.

Since LNAs also have the capacity to bind DNA, there is a possibility that LNA@K1 could interfere with transcription of the miRNA cluster and thus explain such a global effect. We therefore performed RT-qPCR to evaluate the levels of pri-miR-K10/12 in control LNA and LNA@K1 conditions. Figure 4C shows that pri-miR-K10/12 level was not affected by LNA@K1 treatment, ruling out a possible inhibition at the transcription step.

In conclusion, we were able to negatively affect the expression of the 10 clustered miRNAs of KSHV solely by targeting miR-K1 sequence. Since it does not interfere with transcription, this downregulation most likely occurs at the post-transcriptional level, probably by interfering with the Microprocessor recognition and/or cleavage of pre-miR-K1.

### Targeting miR-K1 inhibits the expression of the cluster in infected cells

So far, we have demonstrated that the whole cluster can be downregulated by using one single molecule targeting miR-K1 sequence. However, our experimental settings did not mirror natural conditions of KSHV infection, since the cluster was expressed from a plasmid. In order to test whether this *cis* regulation exists in a context closer to physiological infection, we decided to apply our antisense LNA strategy in HEK293FT cells carrying recombinant KSHV genomes (HEK293FT-rKSHV) (43). Similar to physiological conditions, rKSHV in cells remains mostly in latent state and it produces all viral miRNAs, even though their global expression is much lower, probably due to the number of viral episomes per cell.

By using the same antisense oligonucleotide as previously described (LNA@K1) in these cells, we were not able to see any global downregulation of the cluster, with the exception of miR-K1(data not shown). We hypothesized that most of the LNAs transfected into the cells were probably sponged by the mature miR-K1 molecules already abundantly present in the cell, as opposed to the situation where miRNAs are expressed from a plasmid concomitantly with the LNA treatment. Thus, only a limited amount of the LNA would be available for the microprocessor inhibition and the potential impact on other miRNAs is too low to be measured. To cope with this situation, we designed a new LNA complementary to the opposite strand of pre-miR-K1 stem loop (LNA@K1*). Given that this new molecule will not be sequestrated by the mature miR-K1, more oligonucleotides can reach the nucleus and interfere with pre-miR-K1 processing. In addition, this would also confirm that the effect we observe on the expression of the cluster is dependent on the processing event of pre-miR-K1 and not on a downstream function of the mature miR-K1.

First, to show that LNA@K1* can indeed interfere with processing of miRNA from the cluster, we performed the experiment in HEK293Grip cells by co-transfecting the pri-miR-K10/12 construct with either LNA@K1* or a control LNA and assessed the expression of mature miR-K1, -K4 and -K11 by RT-qPCR. Upon treatment with LNA@K1*, the accumulation of all three miRNAs dropped substantially compared to the treatment with the control LNA, while the level of Let-7a remained unchanged (Figure 5A). As previously, a potential impact of LNA treatment on cluster transcription was verified and only a mild decrease (30%) of pri-miR-K10/12 could be observed between control and LNA@K1* conditions (Figure 5B). We thus confirmed the LNA@K1*-mediated post-transcriptional inhibition occurring during miRNA processing. However, transfection of this LNA oligonucleotide into HEK293FT-rKSHV cells resulted only in a mild impact on KSHV miRNA expression, as demonstrated for miR-K1, -K4 and -K11. While only miR-K11 levels were significantly reduced, miR-K1 and -K4 did not decrease in a statistically significant manner (Figure 5C). This might be explained by a differential stability of the different viral miRNAs in infected cells. In contrast to ectopically expressed cluster, infected cells already contain a certain amount of mature viral miRNAs and their differential turnover would directly influence the sensitivity of our assay. If the half-lives of miR-K1 and - K4 are longer than the half-life of miR-K11, then it might prove difficult to assess the impact of inhibiting their processing. As a solution, we set to measure the accumulation of newly synthesized miRNAs. LNA-transfected cells were therefore incubated with 4-thiouridine (4sU), which allowed the isolation of novel transcripts via selective biotinylation and pull-down on streptavidin beads. Due to the variation between the efficiency of individual pull-downs, viral miRNAs were analyzed in total RNA (input) as well as in isolated pull-down fraction and their enrichment in pull-down over input was expressed relative to Let-7a. Following two different experimental protocols, we observed a consistent and significant reduction in neosynthesized miR-K1, -K3, -K4 and -K11 upon treatment with LNA@K1* (Figure 5D and S8). In addition, we verified by RT-qPCR that the LNA had no impact on the levels of neosynthesized pri-miRNA transcript (Figure 5E). Thus, we have shown that the expression of the KSHV miRNA cluster can be inhibited by using one single oligonucleotide targeting pre-miR-K1 and that this phenomenon indeed takes place also within KSHV-infected cells. Although at this stage, we cannot formally conclude that what is important for the *cis-*regulation is the processing event or the presence of a stem-loop, our results clearly point toward the importance of pre-miR-K1.

**Figure 5.**
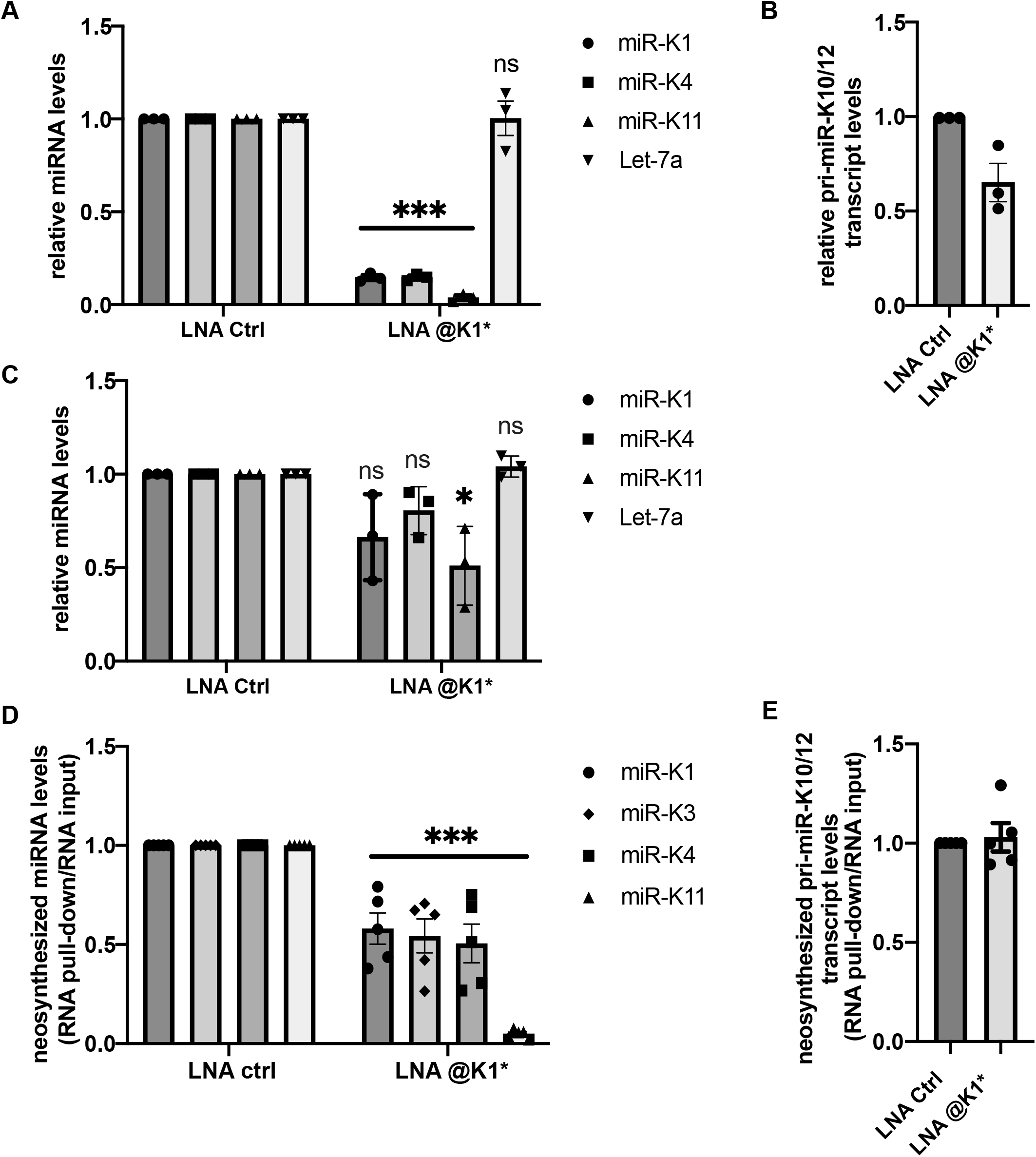
Inhibition of pre-miR-K1 processing impacts the expression of other viral miRNAs in infected cells. (A) Levels of mature miRNAs in HEK293Grip cells co-transfected with pri-miR-K10/12 expression plasmid and LNAs complementary to miR-K1* or control LNA. The analysis was performed on total RNA extracted 48 h post-transfection and miRNA levels were normalized to U48. (B) Measure of pri-miR-K10/12 expression in samples from (A), GAPDH was used as a normalizer. (C) Levels of mature miRNAs in HEK293FT-rKSHVcells transfected twice with 20nM of LNAs complementary to miR-K1* or control LNA. U48 was used as a normalizer. (D) Accumulation of neosynthesized miRNAs in HEK293FT-rKSHV cells transfected with either LNA complementary to miR-K1* or control LNA. 3 h of metabolic labelling with 300 µM 4sU was performed 24 h after LNA transfection and levels of mature miRNAs were measured in total RNA (input) and in isolated newly synthesized fraction (pull-down). To account for variation in pull-down efficiencies, enrichment of miRNA levels in pull-down over input were determined after normalizing to Let-7 levels. (E) Accumulation of neosynthesized pri-miR-K10/12 measured in the samples from (D). The same approach was used to determine the enrichment of pri-miRNA levels in the pull-down over input except that CYC1 was used to normalize the data instead of Let-7. Mature miRNAs and pri-miR-K10/12 in all experiments were quantified by RT-qPCR. All results are displayed relative to control samples set to 1. Bars represent mean ± s.e.m of three (A, B, C) or five (D, E) independent experiments. Statistical significance was verified by unpaired t test with ns: non-significant, *: p < 0.05, ***: p < 0.001.

## DISCUSSION

In this study, we explored a complex layer for miRNA biogenesis regulation in which the expression of miRNA hairpins within a large miRNA cluster is interdependent. We showed that two miRNA hairpins, namely miR-K1 and -K3 hairpins, within the intronic KSHV miRNA cluster were required for the optimal expression of the remaining miRNAs. Indeed, their deletion within an expression plasmid drastically diminished clustered miRNAs expression in cell. Similarly, antisense LNAs that bind to the pre-miR-K1 hairpin and inhibit its processing by impeding the recognition and/or cleavage by the Microprocessor led to global downregulation of the cluster. This strategy also allowed to decrease viral miRNAs in the more natural context of infected cells. Furthermore, our data showed that the pre-miRNA feature *per se*, at least for pre-miR-K1, is responsible for this regulation since pre-miR-K1 could be replaced by a heterologous pre-miRNA. Altogether, these results indicate that miR-K1 and miR-K3 hairpins are important *cis*-regulatory elements for the expression of the KSHV clustered miRNAs. Previously, the Krueger laboratory produced bacmid constructions containing KSHV genome deleted from individual KSHV miRNAs and they also noticed a decrease in expression of other viral miRNAs in mutants deleted of miR-K1 and miR-K3 (50). Here we explain these observations through the *cis-*regulatory function of these two pre-miRNAs.

Interdependency of clustered miRNA hairpins was documented previously for different miRNAs and in diverse species (37–40, 40, 41). However, it has so far only been studied for small clusters of two or three miRNA hairpins where a helper hairpin assists the processing of a neighboring suboptimal hairpin. Here, we show *cis* regulation for the first time within a large cluster of 10 miRNA hairpins. Thus, “assisted” miRNA hairpins are not all proximal to the helper hairpins. In addition, they are not necessarily suboptimal as demonstrated by our 2D structure probing data published previously (31). So, in the case of KSHV miRNAs cluster, *cis* regulation might result from a more complex mechanism. One possibility could rely on the recruitment of the Microprocessor by miR-K1 and miR-K3 hairpins, inducing its local concentration to re-initiate further maturation events on the same polycistronic pri-miRNA.

From a conceptual point of view, this might be compared to ribosome re-initiating translation on a same mRNA. However, from a mechanistic point of view, Microprocessor would not scan the pri-miRNA but rather cycle from the cleaved miRNA hairpin and relocate on promiscuous miRNA hairpin. Two alternate but not mutually exclusive models may help this repositioning. One model would be that the globular and compact 3D structure of pri-miRNA may *per se* help the Microprocessor to relocate. Indeed, structural study performed on pri-miR-17∼92 shows such organization (30). We previously determined the 2D structure of KSHV miRNA cluster using SHAPE method (31). Although we could not conclude on a compact arrangement of the viral pri-miRNA, we did observe numerous stem-loops, containing or not miRNAs, that could participate to long-distance interaction to maintain such compact 3D structure. A second model would involve protein cofactors able to assist in the recruitment of the Microprocessor on the neighboring miRNA hairpin. Very recently, it was shown that ERH and SAFB2 can interact with the Microprocessor and their ability to dimerize may even mediate multimerization of the Microprocessor and allow its simultaneous binding to several hairpins (37, 40, 42, 51). However, this model remains to be clearly established. For example, the role of ERH and SAFB2 proteins may be specific to only certain miRNA clusters and not all clustered miRNAs may depend on such assistance. Indeed, using CRISPR/Cas9 editing, Lataniotis et al showed that editing of miR-195 led to a decrease of its neighboring miR-497, whereas no such interdependency was observed for miR-106∼25 or miR-17∼92 clusters (39). Another study, based on genetically engineered mice, also showed that deletion of any miRNA from the cluster miR-17∼92 did not alter the expression of the other miRNAs (52). In the case of the KSHV cluster, pre-miR-K1 and pre-miR-K3 may interact with high affinity with a protein cofactor, potentializing further interaction with the other pre-miRNAs. This protein cofactor might then recruit or improve the Microprocessor activity.

Another mechanism may rely on specific structural constrains that could be resolved after cleavage of miR-K1 and miR-K3 hairpins, rendering the other miRNA hairpins more accessible to the Microprocessor. Indeed, previous studies reported that the globular fold of the pri-miR-17∼92 may autoregulate its processing (30, 53, 54). Chaulk et al. demonstrated that the compact fold of the cluster is adopted through a specific tertiary contact between pre-miR-19b and a non-miRNA hairpin resulting in reduced miR-92a expression whereas this inhibitory effect could be abolished by disrupting this contact (54). However, our data obtained from our *in vitro* processing assays of RNA mimicking initial cleavage of miR-K1 or miR-K3 hairpins show neither global nor drastic improvement of other miRNA hairpins maturation. This suggests that miR-K1- and miR-K3-dependent regulation may rather occur *in cis*, when the two hairpins are still present in the cluster. Cleavage of the cluster may also happen in a hierarchical way similarly to pri-miR-17∼92 (55). In that case, the restricted impact of pre-miR-K1 or pre-miR-K3 cleavage may reflect that processing of the KSHV cluster occurs in different steps having each downstream additive positive effect. Unfortunately, the experimental design of our *in vitro* processing assays did not allow to follow complete processing of the cluster. Thus, we probably observed only the first cleavage events.

Another intriguing aspect of this work is the fact that despite their efficient processing by the Microprocessor, the levels of mature miR-K1 and -K3 are low in infected cells. It would be interesting to know if these two pre-miRNAs are as well efficiently exported into the cytoplasm and/or processed by Dicer and how the excess of pre-miRNAs is eliminated from the cell. It was shown previously that MCP-1-induced protein-1 (MCPIP1), a suppressor of miRNA biogenesis and involved in immunity, could directly cleave KSHV pre-miRNAs through its RNase domain (56, 57). However, while their results show that pre-miR-K1 and -K3 can be bound and cleaved by MCPIP1, almost all remaining KSHV pre-miRNAs are prone to be degraded as well, suggesting that another protein or mechanism might be involved in selective decay of excessive pre-miR-K1 and –K3.

KSHV miRNAs are involved in a multitude of functions related to the viral life cycle, immune escape, and pathogenesis (reviewed by e.g. (58, 59)). Latent infection is one of the biggest hurdles in the treatment of KSHV-related pathologies. Therefore, the role of viral miRNAs in latency maintenance is of great importance. Several groups have shown that artificial reduction of levels of particular KSHV miRNAs can lead to higher viral reactivation. For example, the main transactivator of lytic cycle RTA is directly regulated by at least two miRNAs, miR-K7-5p and miR-K5 (10, 60). In addition, miR-K1 indirectly controls latency maintenance by downregulation of the inhibitor of NF-kB pathway, IkBa (9). Furthermore, many of their protein targets validated to date are involved in pathways important in oncogenesis (61). Interestingly, miR-K11 presents the same seed sequence as a well-known oncomiR miR-155 and both miRNAs target a common subset of genes (62, 63). The Gao’s and Renne’s groups have shown that several KSHV miRNAs participate to the transforming potential of KHSV by targeting cell growth and survival pathways (64, 65). Targeting several, if not all the KSHV miRNAs can therefore represent a valuable therapeutic option. Recently, Ju et al. have proposed a therapeutic strategy based on LNA-modified oligonucleotides complementary to miR-K1, - K4 and –K11 coupled to carbon dots for better intracellular delivery (66). While they use a combination of three different molecules, our results suggest that blocking the processing of one single miRNA might lead to a global decrease of the entire miRNA cluster. Given that about 25 to 40% of all human miRNAs are embedded in clusters (25, 26), and given the growing body of evidence that they are implicated in disease, the ability to suppress their expression as a whole might be of importance for future therapies.

## SUPPLEMENTAL MATERIAL

Supplemental material is available for this article.

## AUTHOR CONTRIBUTIONS

AF, SP, MC and MV conceived the project. MC, MV, SP and AF designed the work. MC, MV, RR and AF performed the experiments and analyzed the results. PD performed the kinetic analysis. EE and PMO generated the HEK293FT-rKSHV cells. AF and SP coordinated the work and SP assured funding. MV, PD, SP and AF wrote the manuscript with input from the other authors. All authors reviewed and approved the final manuscript.

## ACKNOWLEDGEMENTS

The authors would like to thank members of the Pfeffer laboratory for discussion. They also would like to thank Prof Narry Kim for the kind gift of plasmids expressing flag-tagged Drosha and DGCR8.

## SUPPLEMENTAL MATERIAL

### SUPPLEMENTAL METHOD

#### Kinetic analysis

##### Practical problems and fitting method

There is a significant upward curvature of the low-amplitude cleavage curves around t = 0 (see for example pre-mir-K2 and pre-miR-K7 for 20130530). This was interpreted as a linearity problem of the IP response since this is only visible for the low-amplitude curves (accordingly, it is almost invisible with the Exp#1 data having higher amplitudes). Such a feature cannot be accounted for by equations (3) and (3’) (see main manuscript) imposing a downward curvature around t = 0. In order to nevertheless use these equations, a simple correction was devised to mimic this lack of linearity for the lowest values of *K*_*i*_(*t*)/*R*_0_. For this, instead of using directly *Y*_*i*_ = *K*_*i*_(*t*)/*R*_0_ to fit the experimental curves, a modified value of *Y*_*i*_ was used according to the response function:

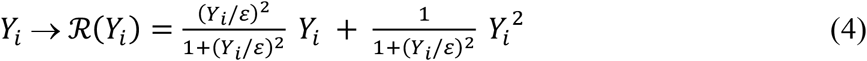

with *ε* a small value. This response function gives 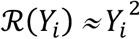 for *Y*_*i*_ of order *ϵ*, which provides the upward curvature close to *Y*_*i*_ = 0, and it is transformed smoothly into ℛ(*Y*_*i*_) = *Y*_*i*_ for increasing values of *Y*_*i*_ above *ϵ*. This *ad-hoc* procedure was quite effective to improve the quality of results, particularly for the Exp#2 data. The value of *ε* was tuned to 0.5 % to obtain the best fit of all curves with a minimum value of the global sum of the errors on *f*_*i*_ and 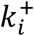 for both data sets. Such a low value indicates that the correction is indeed a minor one. Practically, the fit was done with the function NonlinearModelFit (with Method → “ConjugateGradient”) in *Mathematica* V.11 from Wolfram Research.

##### Using a more stringent kinetic model of pri-miRNA cleavage

The simple model used in the study allowed us to obtain excellent fits (Figure 1 and Figure S1), but with three free parameters (*f*_*i*_, 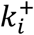 and 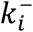) per experimental curve. In order to use a more stringent test, we imposed two restrictions for a better representation of reality. First, we imposed that the variations of the rates of cleavage by Drosha from one experiment to another one should only result from the amount of Drosha in each experiment. For this, we imposed a strict proportionality of the two sets of 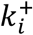. Second, we imposed that the cleaved fraction *f*_*i*_ of *R*_0_ yielding the pre-miR *K*_*i*_ was the same for the two experiments (Figure S2). This more stringent method, therefore, involves only two adjustable parameters per curve, which represents quite a significant reduction of the degrees of freedom.

The fitting of the experimental cleavage curves with the kinetic model with three free parameters (see Material and Methods) per curve led to excellent agreement (Figure 1B and Figure S1B). Note that the slight correction for non-linearity (see Supplemental Material) was important to obtain this result. As expected, the results with a more stringent model with only two free parameters per curve were less good but mostly for the low-amplitude curves with the lowest signal-to-noise ratio (Figure S2). This indicates that the excellent agreement with three parameters was not simply the result of a meaningless numerical fit, which is in good support for the simple kinetic model in use. The numerical results are shown in Table S1.

##### *In vitro* pre-miRNA stability assays

Measure of stability of *in vitro* transcribed pre-miRNAs was performed in the same conditions as the in vitro Drosha miRNA processing assays (see Materials and Methods) except that whole cell lysate was used without overexpressing Drosha and DGCR8. Briefly, 500 fmol of each of the four tested pre-miRNAs were pooled together, denatured, let to refold and incubated in total HEK293Grip cells extract for increasing times at 37°C. 1/10 of the phenol-extracted and ethanol-precipitated RNAs was used for northern blot analysis. A standard curve was generated by loading decreasing amounts (50 to 3.125 fmol) of corresponding *in vitro* transcribed pre-miRNAs.

## SUPPLEMENTAL TABLES

**Table S1.**
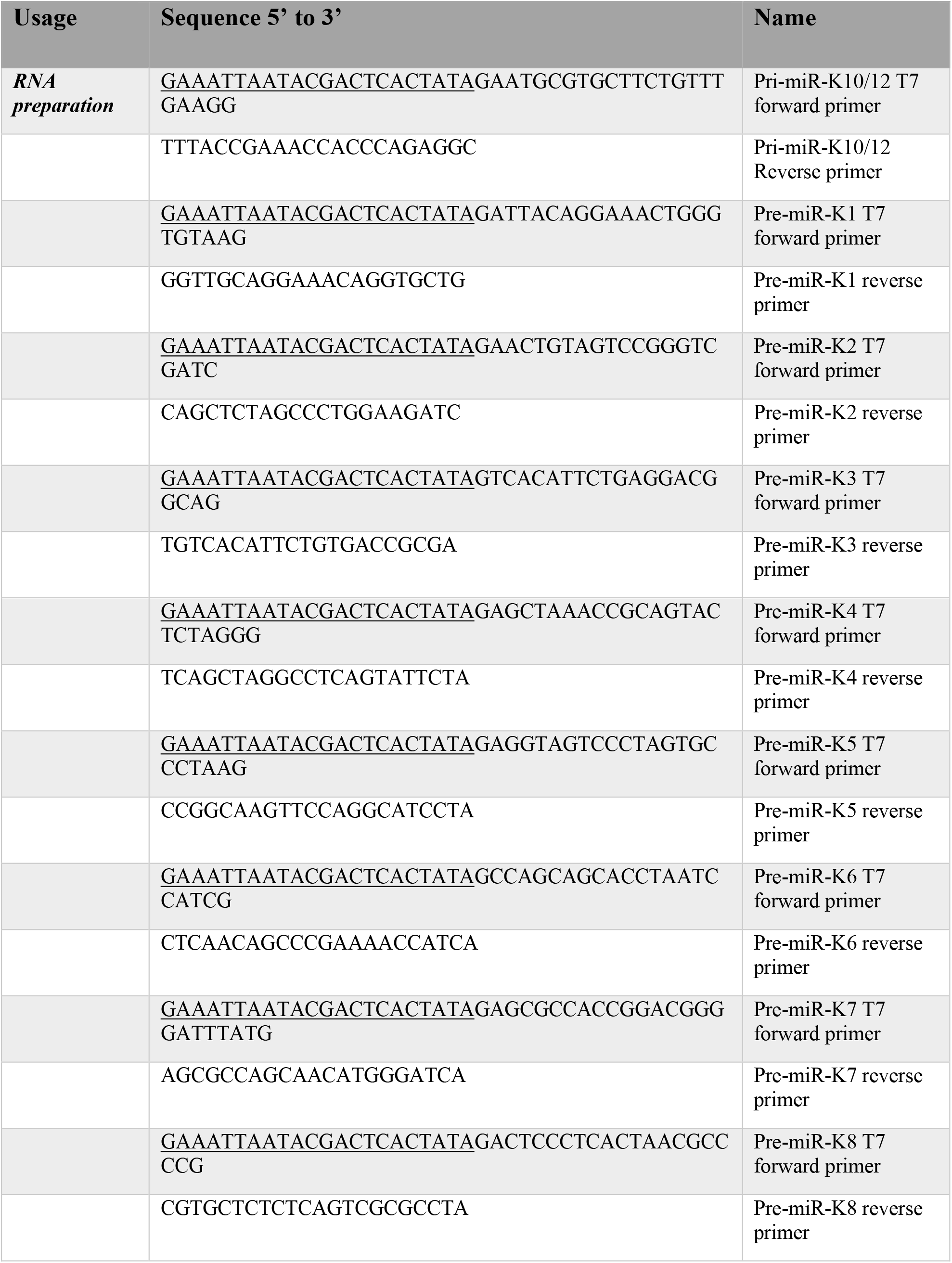
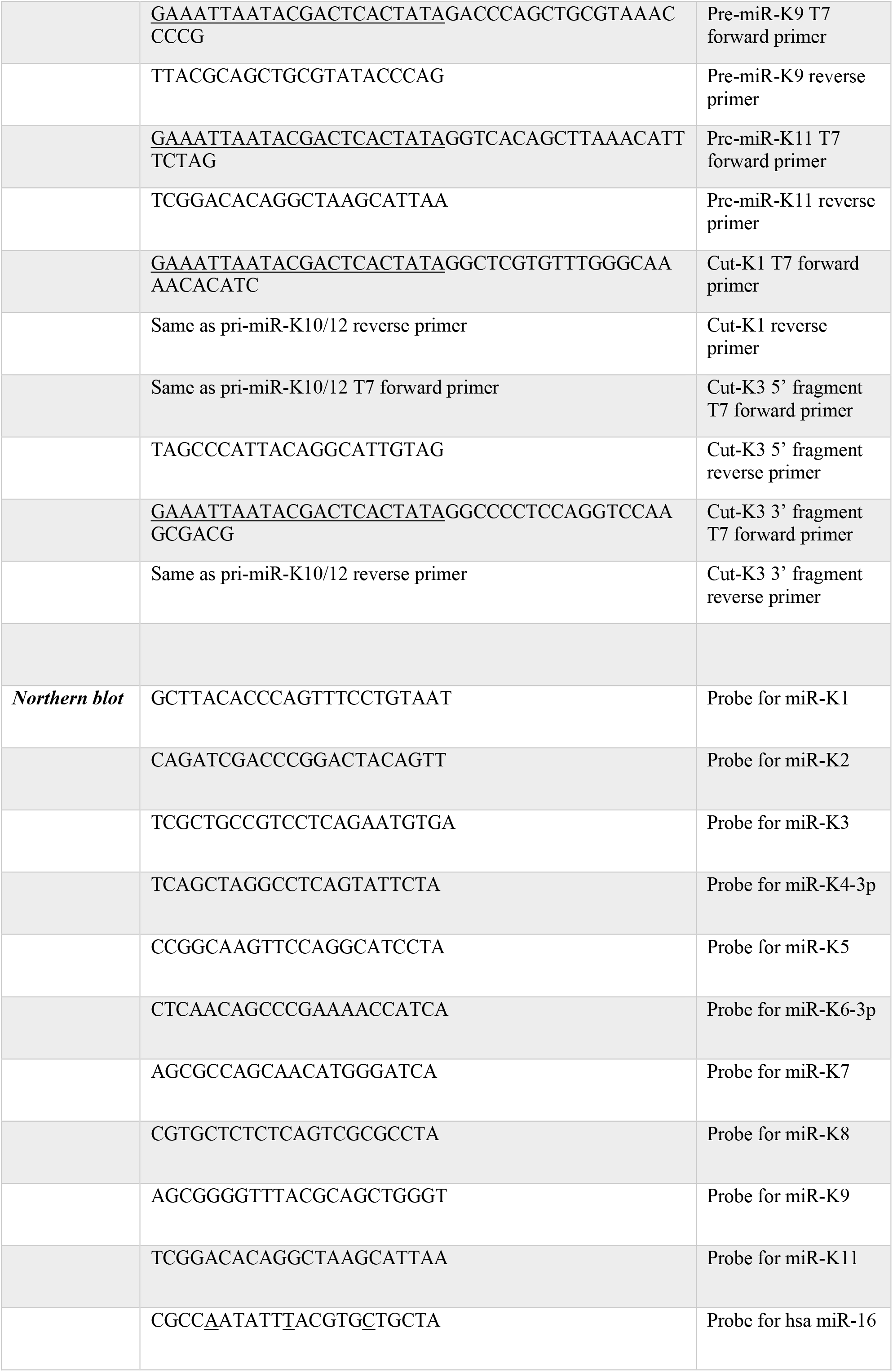
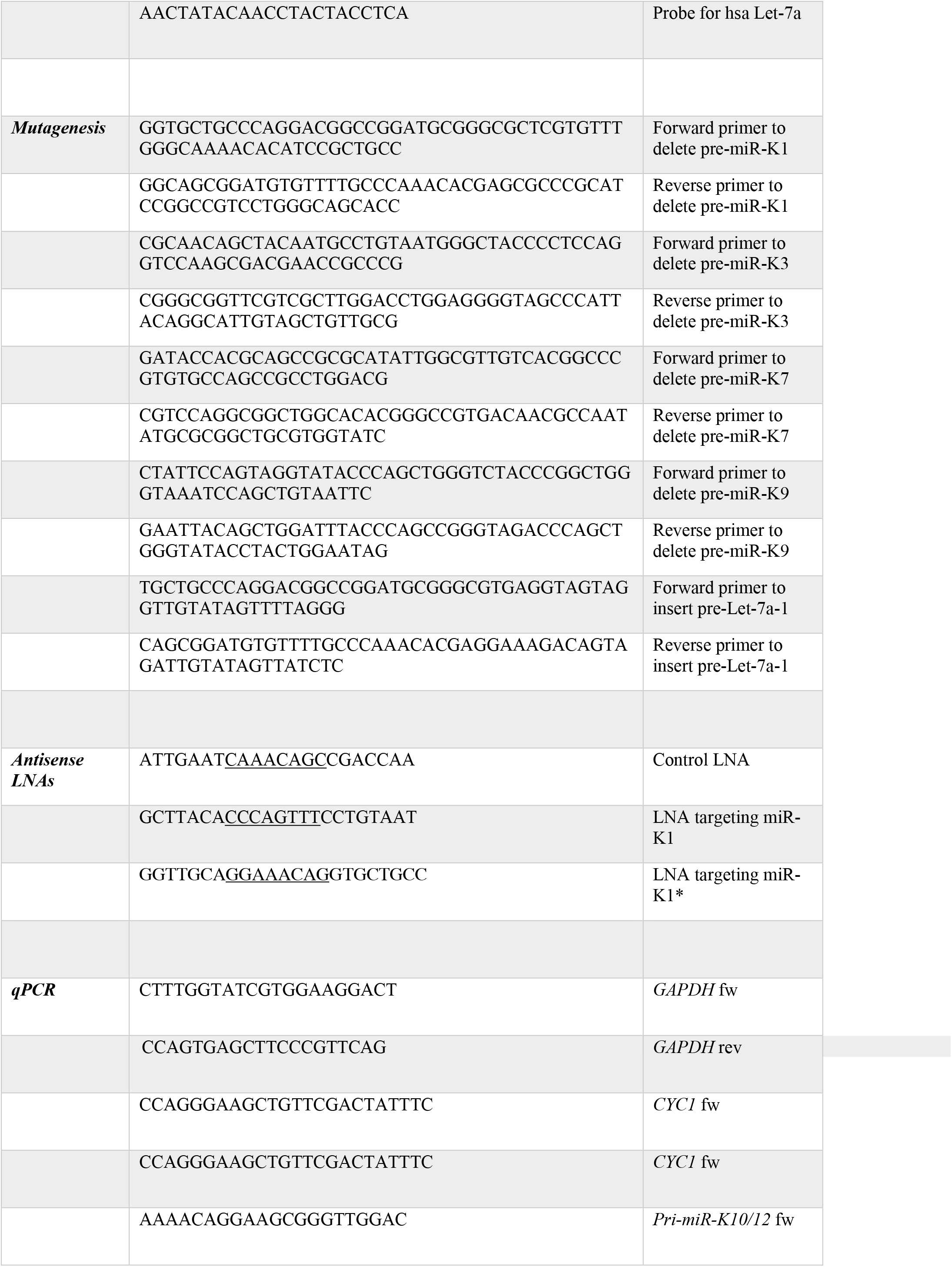
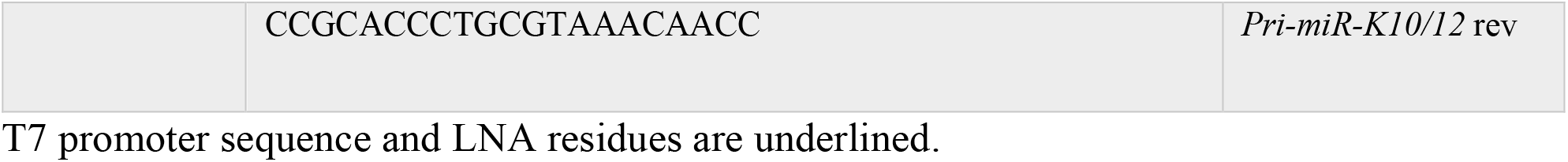
Sequences of oligonucleotides used in this study.

**Table S2.** Kinetic parameters for cleavage of KSHV miRNA hairpins within pri-miR-K10/12 by the Microprocessor *in vitro*.

| pre-miR | Exp#1 |  |  | Exp#2 |  |  |
| --- | --- | --- | --- | --- | --- | --- |
| | $f(\%)$ | $10 \times k^+$<br>(min<sup>-1</sup>) | $100 \times k^-$<br>(min<sup>-1</sup>) | $f(\%)$ | $10 \times k^+$<br>(min<sup>-1</sup>) | $100 \times k^-$<br>(min<sup>-1</sup>) |
| **K1** | 15.0 ± 1.0 | 1.50 ± 0.20 | 1.10 ± 0.20 | 17.0 ± 0.6 | 0.30 ± 0.01 | 3.00 ± 0.20 |
| **K2** | 1.7 ± 0.2 | 0.97 ± 0.20 | 0.32 ± 0.30 | 1.0 ± 0.1 | 0.53 ± 0.08 | 0.80 ± 0.30 |
| **K3** | 32.0 ± 2.0 | 0.82 ± 0.08 | 0.34 ± 0.20 | 40.0 ± 0.9 | 0.22 ± 0.005 | 2.20 ± 0.10 |
| **K4** | 15.0 ± 5.0 | 0.46 ± 0.20 | 1.70 ± 1.00 | 12.0 ± 12.0* | 0.18 ± 0.18* | 1.80 ± 1.80* |
| **K5** | 11.0 ± 0.8 | 0.81 ± 0.09 | 0.57 ± 0.20 | 2.8 ± 0.6 | 0.58 ± 0.20 | 0.15 ± 0.50 |
| **K6** | 6.1 ± 5.0 | 0.38 ± 0.40 | 0.00 ± 2.00 | 2.8 ± 2.0 | 0.41 ± 0.30 | 0.00 ± 1.00 |
| **K11** | 11.0 ± 0.3 | 1.60 ± 0.10 | 1.10 ± 0.10 | 9.6 ± 9.6* | 0.26 ± 0.26* | 2.60 ± 2.60* |
| **K7** | 3.4 ± 0.06 | 0.27 ± 0.005 | 2.70 ± 0.10 | 3.7 ± 0.1 | 0.14 ± 0.004 | 1.40 ± 0.10 |
| **K8** | 11.0 ± 0.5 | 2.30 ± 0.30 | 0.10 ± 0.10 | 13.0 ± 3.0 | 0.71 ± 0.20 | 1.80 ± 0.90 |
| **K9** | 7.0 ± 1.0 | 0.56 ± 0.10 | 0.05 ± 0.50 | 7.7 ± 6.0 | 0.32 ± 0.30 | 0.78 ± 2.00 |

Pre-miRNA accumulation levels $f$ are in percentage of cleaved pri-miRNA in the assay (initial concentration = 16.7 nM, see Material and Methods). Cleavage rate constants $k^+$ and $k^-$ are associated to cleavage by the Microprocessor or to residual Dicer or another RNase activity, respectively.
\* when errors were extremely higher than the determined values, they were arbitrarily set at 100 %.

**Table S3.**
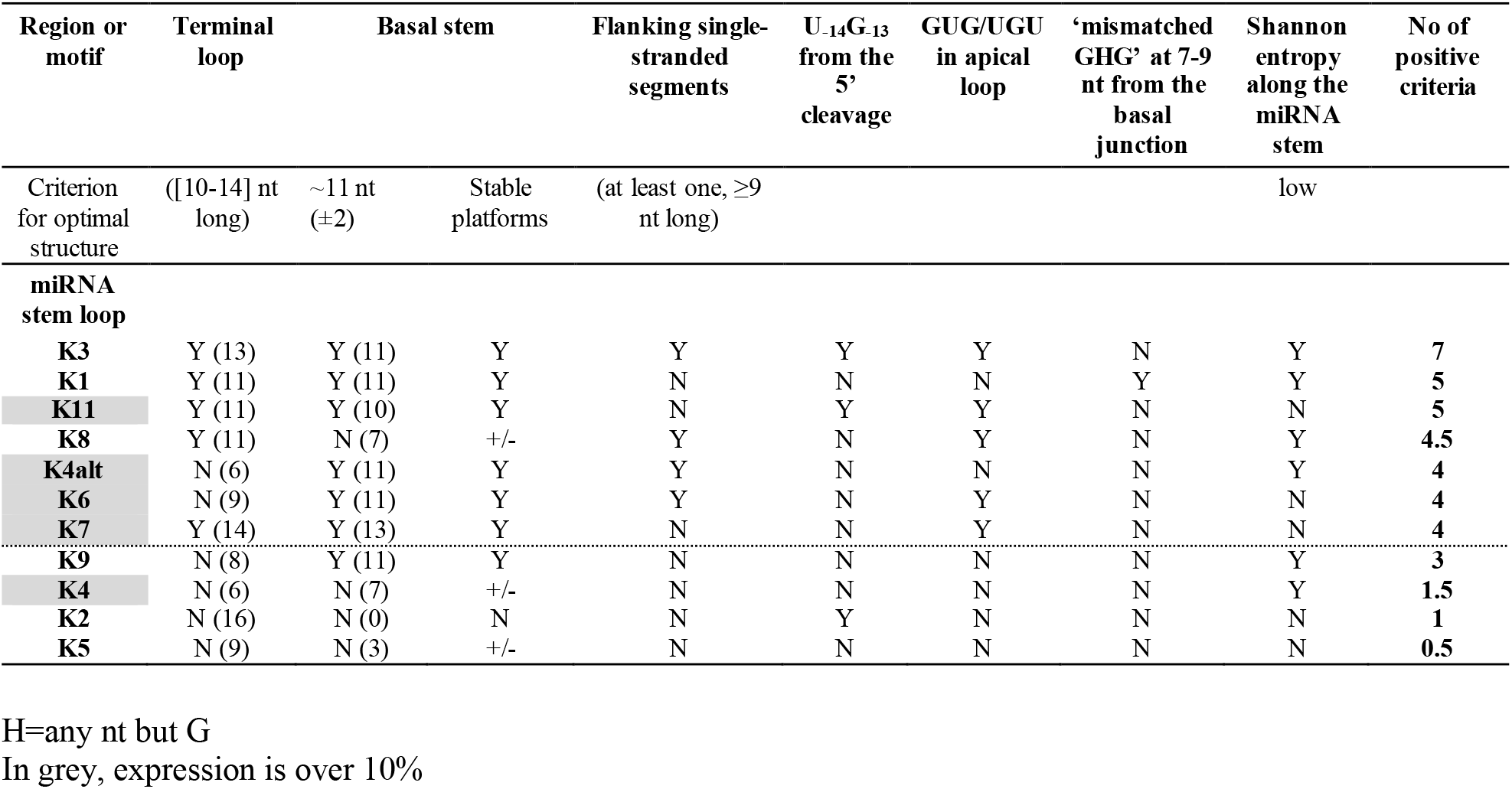
Primary sequence and structural features determinants of miRNA stem-loops from KSHV pri-miR-K10/12.

## SUPPLEMENTAL FIGURES AND LEGENDS

**Figure S1.**
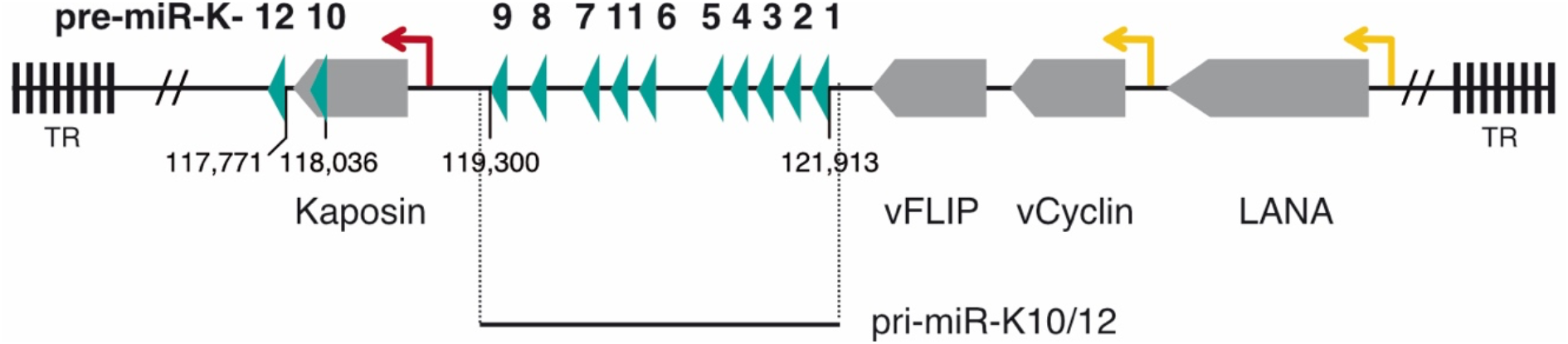
Genomic organization of KSHV miRNAs and location of pri-miR-K10/12. The twelve KSHV pre-miRNAs are localized in the latency locus and are indicated by green arrow heads. Ten of them are clustered in an intron (pre-miR-K1 to -K9 and pre-miR-K11) from which the sequence referred to as pri-miR-K10/12 derives, whereas pre-miR-K10 and - K12 are in the coding region and in the 3’UTR of Kaposin mRNA, respectively. Sequence coordinates were derived from reference sequence NC_009333.1. Open reading frames are in grey. Lytic promoter is represented by a red arrow and latent promoters by yellow arrows. TR, Terminal Repeats.

**Figure S2.**
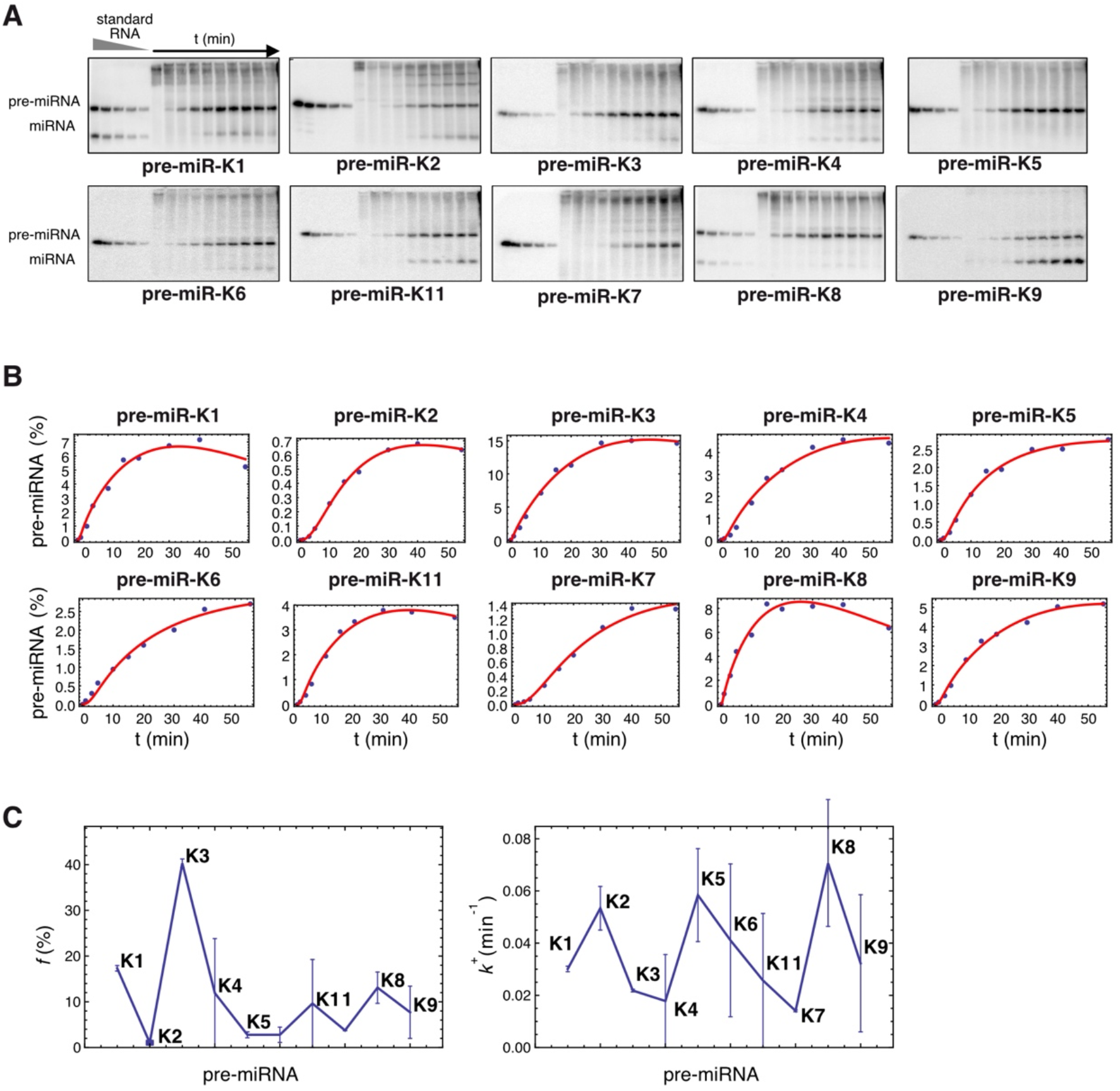
Kinetic analysis of KSHV clustered pre-miRNAs maturation *in vitro* by the Microprocessor (Exp#2). (A) Northern blot analysis of the time course of *in vitro* processing assays using *in vitro* transcribed pri-miR-K10/12 and Hek293Grip cells total protein extract where Drosha and DGCR8 were overexpressed. *In vitro* transcribed pre-miRNAs and synthetic RNA oligonucleotides were loaded at decreasing concentrations as standards. (B) Cleavage curves were obtained after plotting pre-miRNA product, in percentage of initial pri-miR-K10/12 substrate, according to time. The fits were obtained with the model involving three free parameters per curve (compare with Figure S3 for the more stringent model with two free parameters per curve). (C) Processing efficiencies (left panel) and cleavage rate (right panel) were plotted in respect to miRNA hairpins showing variation among the clustered pre-miRNAs. The error bars come from standard procedures used to fit the experimental curves by minimizing the residuals between the experimental points and their theoretical estimates.

**Figure S3.**
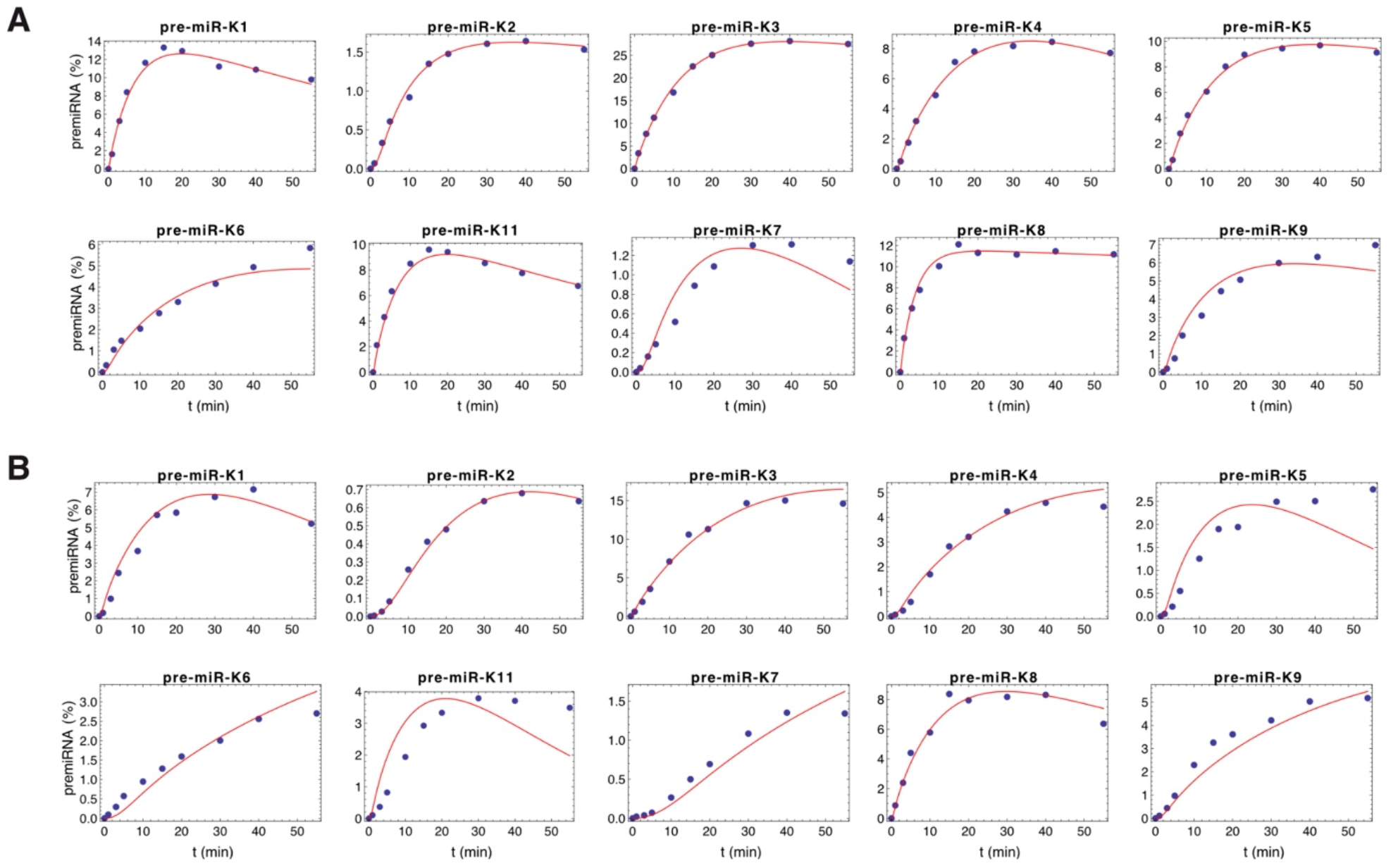
Joint fitting of the experimental curves for Exp#1 (A) and Exp#2 (B) with two free parameters per curve.

**Figure S4.**
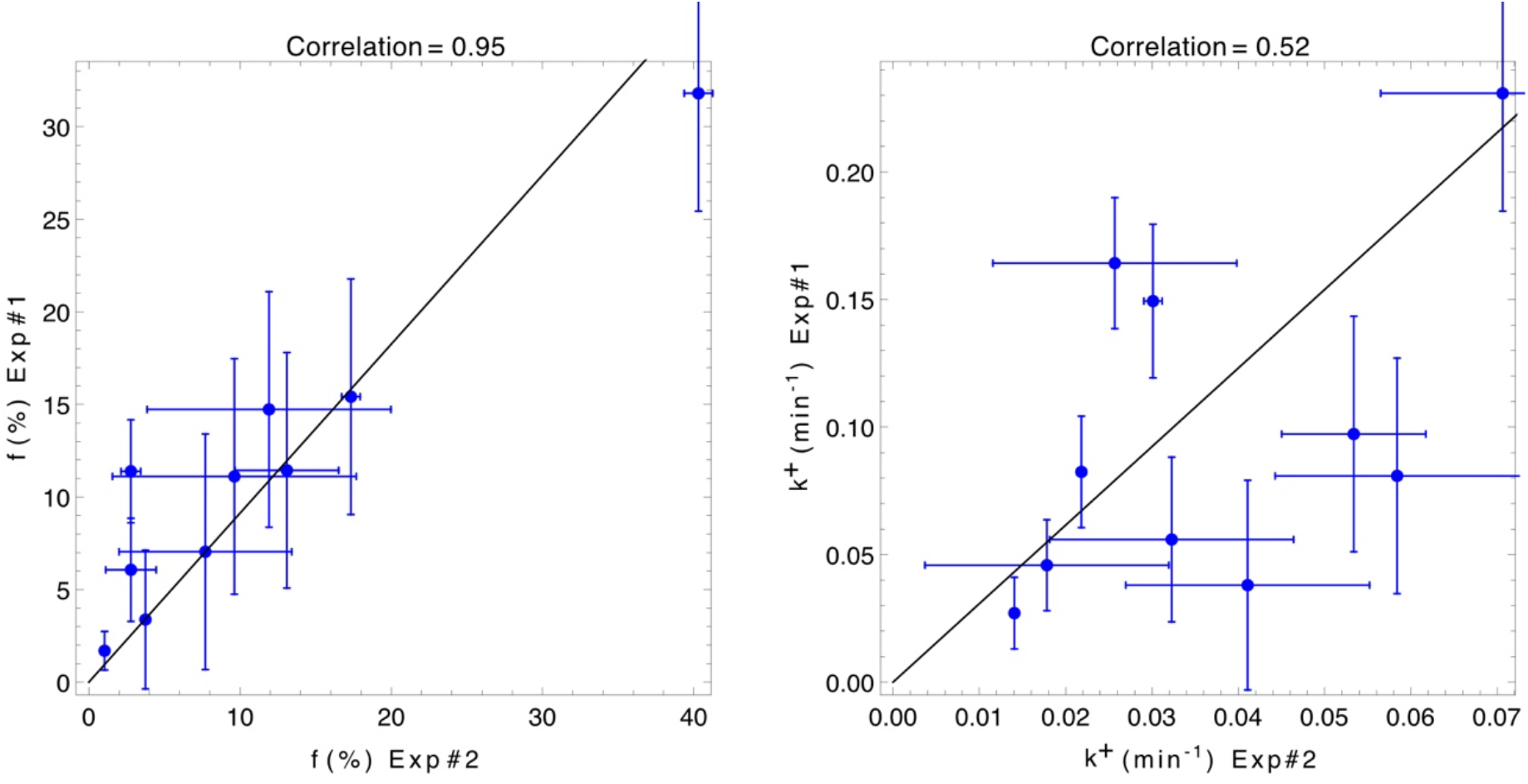
Correlation among experiments of *in vitro* processing assays. Processing efficiencies (*f* in percentage, left panel) and cleavage rate constants (*k*^+^ in min^-1^, right panel) were compared between the two experiments analyzed in this study, namely Exp#1 and Exp#2.

**Figure S5.**
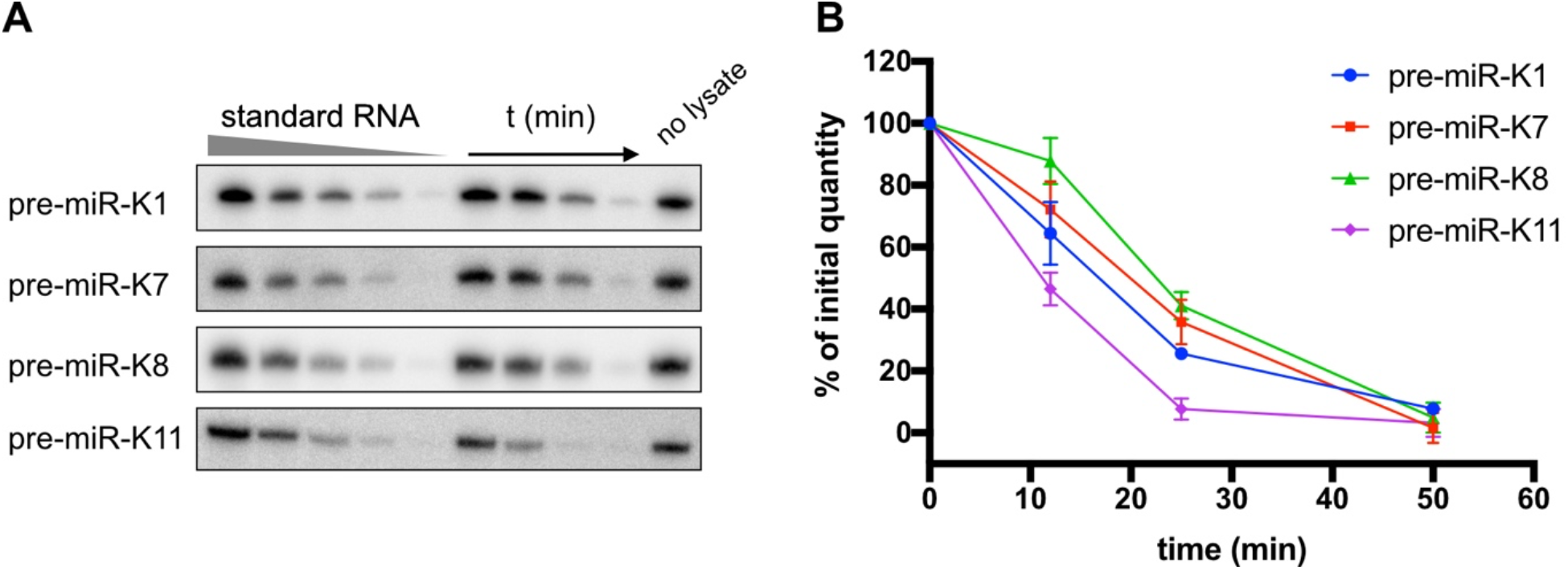
Determination of synthetic pre-miRNAs stability in processing assays. *In vitro* transcribed pre-miRNAs were incubated in whole cell lysate from HEK293Grip cells and submitted to conditions used for *in vitro* processing assays. Their decay was followed over time. Northern blots (A) were quantified by using standard pre-miRNAs and results from three replicates were plotted (B) relative to pre-miRNA quantity at 0 min.

**Figure S6.**
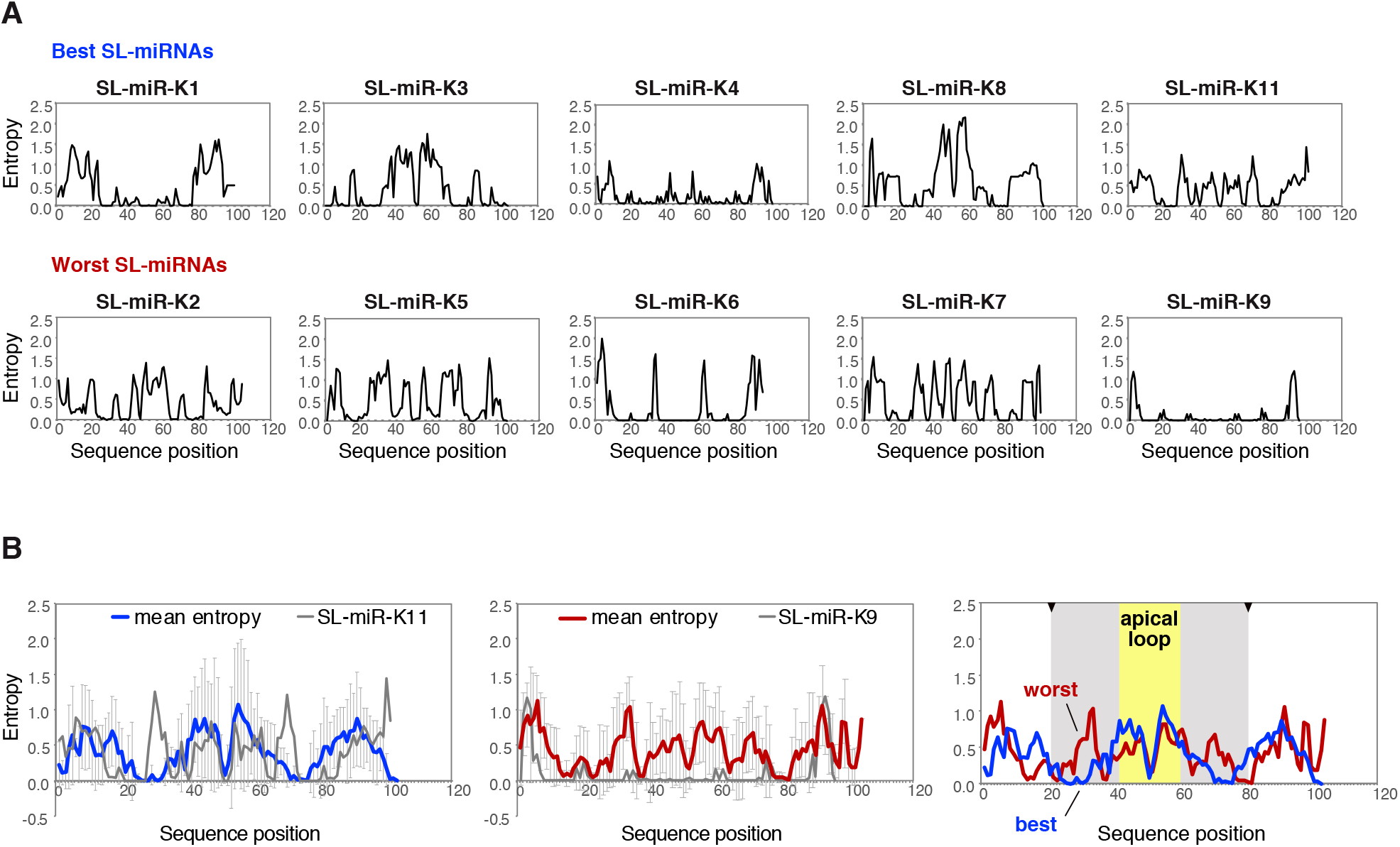
Positional entropy of KSHV miRNA hairpins. (A) Shannon entropy was plotted across the sequence of individual miRNA hairpins, namely stem-loop (SL)-miRNAs, including the pre-miRNA plus 20 nucleotides on both sides, using RNAfold from ViennaRNA Web Services (Institute for Theoretical Chemistry, University of Vienna) (1, 2). (B) Mean entropies of the best substrates (processed over 10%, blue curve, left panel) and of the worst substrates (processed below 10%, red curve, middle panel) were plotted across the sequence. Comparison of the two curves (right panel) show that the best substrates are enriched for low entropy along the stem in contrast to the worst substrates, in agreement with data published in Rice et al (3). However, in the two groups, exceptions come with SL-miR-K11 (processed over 10% but showing high entropy along the stem, grey curve, left panel) and SL-miR-K9 (processed below 10% but showing low entropy, grey curve, middle panel). The approximate location of apical loop and the stem is highlighted in yellow and grey, respectively, and cleavage sites are indicated by arrow heads.

**Figure S7.**
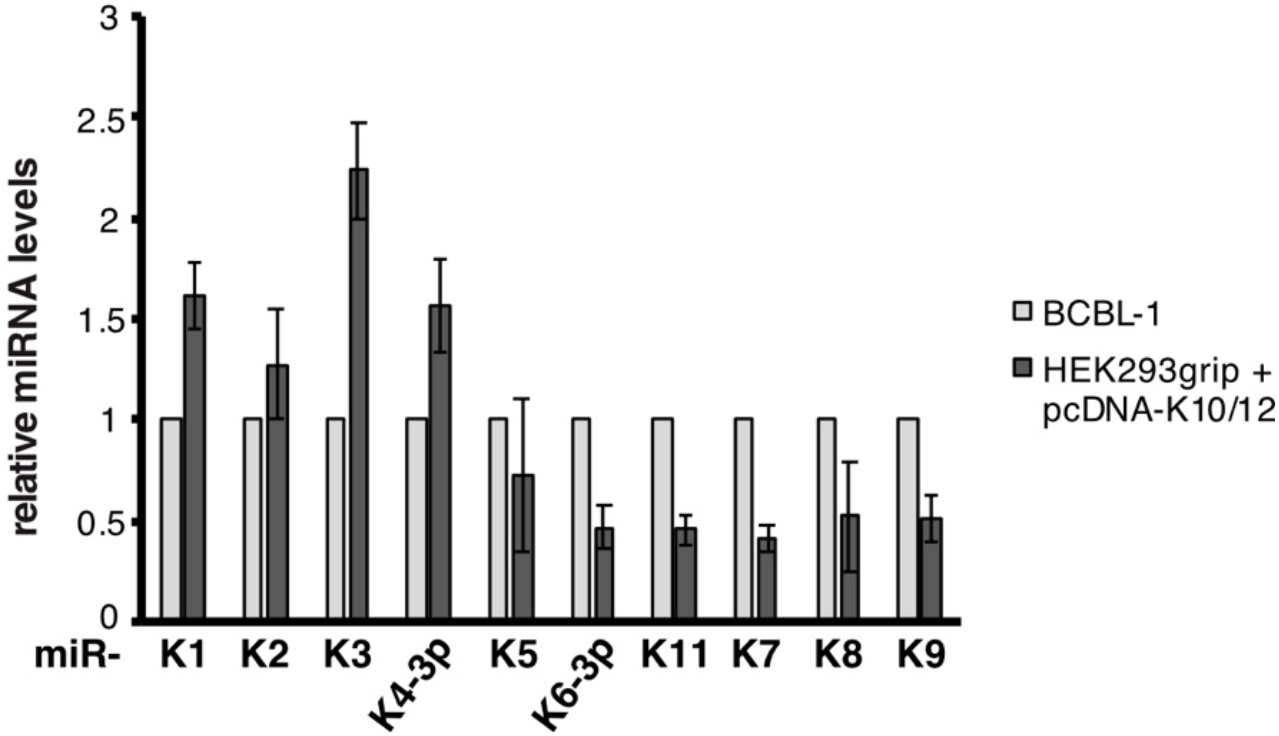
Relative expression of KSHV miRNAs from HEK293grip cells transiently transfected with pcDNA-K10/12 compared to expression in BCBL-1 infected cells. Values were obtained by quantifying signals from northern blot analysis. Error bars derive from three independent experiments except for miR-K1, -K2, -K4-3p and -K6-3p where n=2.

**Figure S8.**
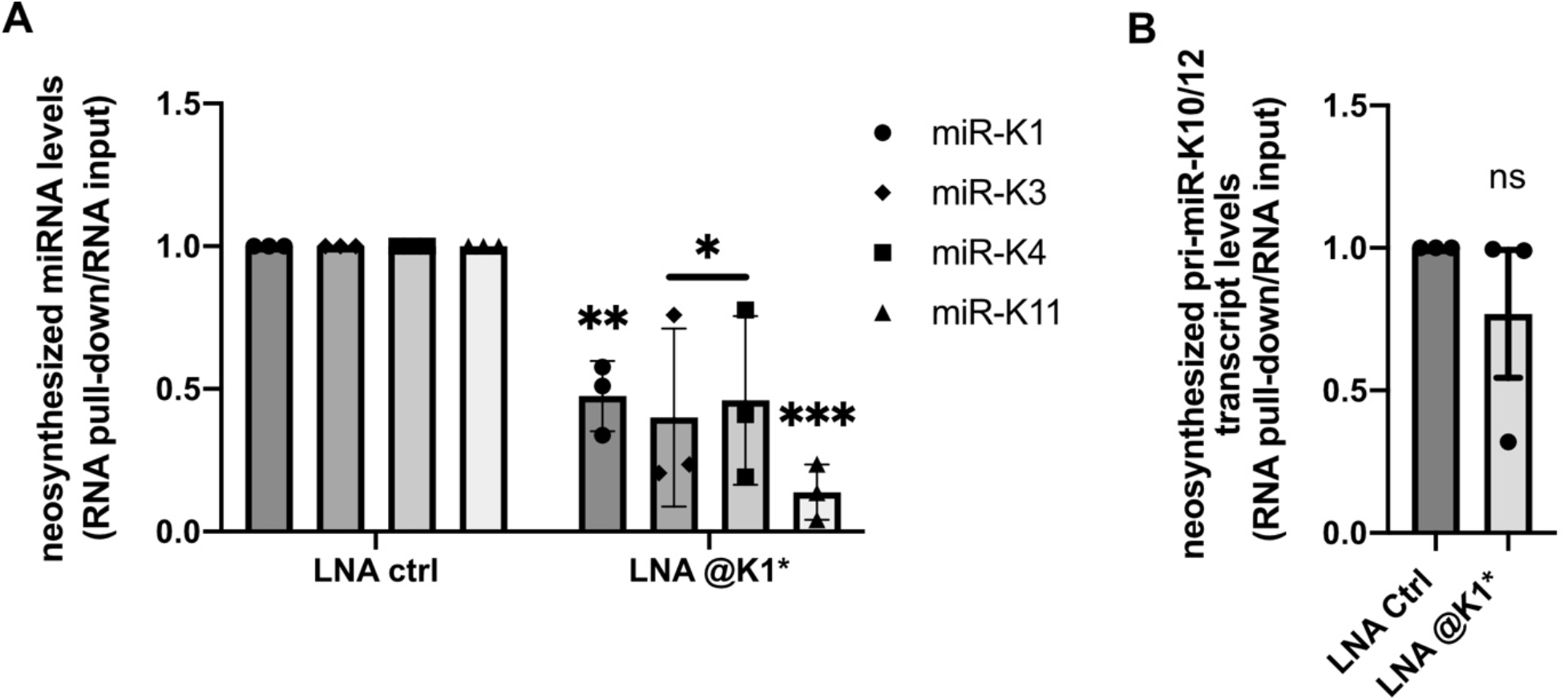
Quantification of neosynthesized miRNAs upon treatment with 20 nM of LNA oligonucleotides. HEK293FT-rKSHV cells were transfected with 20 nM LNA complementary to miR-K1* or control LNA. 24 hours after transfection, they were incubated with 100µM 4sU for another 16 hours. Neosynthesized transcripts having incorporated 4sU were isolated and levels of mature miRNAs (A) and primary transcript (B) were measured by RT-qPCR. Histograms show ratios of enrichment in pull-down over input RNA relative to Let-7 levels which were set to 1 in control samples. Enrichment of primary transcript was determined relative to CYC1. Bars represent mean ± s.e.m of three experiments. Statistical significance was verified by unpaired t test with ns: non-significant, *: p < 0.05, **: p < 0.01, ***: p < 0.001.

